# Scalable multimodal mapping of macrophage regulatory architecture by integrating optical and transcriptomic pooled screens

**DOI:** 10.64898/2026.05.27.728345

**Authors:** Takamasa Kudo, Romain Lopez, Ana M. Meireles, Antonio Rios, Paula Coelho, Vineethkrishna Chandrasekar, Dylan C. Lam, Jan-Christian Huetter, Mineto Ota, Orit Rozenblatt-Rosen, Levi Garraway, Kathryn Geiger-Shuller, Avtar Singh, Jonathan K. Pritchard, Aviv Regev

## Abstract

Understanding how genetic perturbations reshape cellular states requires measuring diverse phenotypic modalities at scale. Here, we present PerturbPair, a platform that combines parallel Perturb-Seq and optical pooled screening (OPS) in primary mouse bone marrow-derived macrophages stimulated with lipopolysaccharide. Profiling over 334,000 single-cell transcriptomes across ∼1,000 gene perturbations and 7.8 million imaging-phenotyped cells across ∼3,000 gene perturbations, we reveal high concordance between transcriptomic and optical perturbation signatures. While RNA and imaging phenotypes were broadly concordant, OPS demonstrated superior sensitivity for weak-effect perturbations owing to greater cell throughput, and captured post-transcriptional regulatory events—such as protein trapping and mTOR-dependent phosphorylation—that left no detectable transcriptional footprint. To exploit cross-modal relationships, we developed EB-MoCAVI, an empirical Bayesian variational inference framework that both imputes RNA profiles for perturbations measured only by imaging—effectively tripling our Perturb-Seq dataset *in silico*—and denoises transcriptomic estimates for perturbations with sparse cell coverage by leveraging matched imaging data as a regularizer. A secondary screen validated the imputed transcriptional profiles and corroborated the cytosolic iron-sulfur assembly pathway as a candidate restraint on macrophage interferon tone. Integrating measured and imputed profiles with rare-variant burden statistics from the UK Biobank identified disease-specific macrophage gene programs and causal regulatory nodes for monocyte counts, type 2 diabetes, and inflammatory bowel disease. PerturbPair establishes a generalizable framework for multimodal perturbation atlases, pointing toward quantitative, causally-informed cross-modal models of cellular behavior.

## INTRODUCTION

Understanding how cells respond to genetic perturbations is fundamental to deciphering the causal mechanisms that govern cellular behavior^1^. The introduction of pooled screens with high-content readouts over the past decade is set to revolutionize our ability to decipher regulatory circuits at scale. Perturb-Seq combines CRIS-PR-based genetic perturbations with single-cell genomics readouts, such as single-cell RNA-seq (scRNA-seq), allowing researchers to profile in a single pooled experiment the impact of thousands of perturbations^2–4^ on rich, interpretable molecular readouts, such as gene expression profiles, protein abundance^5–8^, and/or chromatin accessibility^9–11^. A complementary approach, optical pooled screening (OPS), couples pooled genetic screens to high-content imaging readouts^12–14^. In OPS, cells under different CRISPR perturbations are identified via optical barcodes or in situ sequencing (ISS), such that microscopy captures both morphological phenotypes and perturbation identity at a massive scale. While Perturb-Seq excels at revealing molecular programs, OPS can capture cell biology features such as organelle organization, protein localization, and dynamic behaviors in live cells. Across both strategies, systematically perturbing genes and measuring complex cellular responses across multiple features has been used for mapping regulatory networks that control cell fate^15^, function^2,16,17^, and dysfunction in disease^6,11,18,19^.

A key limitation remains: although we can now measure diverse classes of phenotypes—such as RNA profiles, protein levels, and morphological features—our understanding of how perturbations affect the cell as a whole remains very limited^20^. Systematically mapping these cross-modal relationships at scale promises to characterize how molecular and morphological phenotypes are co-regulated, an essential stepping stone towards building comprehensive models of cell biology^1,21^, sometimes called “virtual cells”^22^. Such models can reveal how different organizational levels—transcriptomic, proteomic, and structural—relate to and influence one another, highlighting where these layers are concordant or divergent. However, doing so requires new experimental assays that produce matched multi-modal perturbation data as well as computational frameworks that integrate across heterogeneous modalities. Additionally, as a practical consequence, if one “cheaper” modality (e.g., imaging) could reliably predict another (e.g., RNA), it would dramatically reduce experimental costs and complexity: enabling researchers to perform inexpensive, large-scale imaging screens to impute costly molecular readouts^1^.

Here, we present PerturbPair—an integrated platform for matched (parallel) Perturb-Seq and OPS with novel AI algorithms that can integrate both screens and predict perturbation effects in one modality from the other. We used a pooled CRISPR library in primary mouse bone marrow-derived macrophages (BMDMs) stimulated with lipopolysaccharide (LPS), a model system characterized by dramatic expression changes and striking morphological transformations. In this system, we captured in two parallel experiments each perturbation’s effect on gene expression (by Perturb-Seq) and cellular morphology (by PerturbView^14^, a scalable and efficient OPS method). We devised analytical approaches to directly compare RNA and imaging phenotypes at scale, revealing which cellular processes are better captured by each modality and identifying perturbations that would be missed by either approach alone. We also developed an AI framework, EB-MoCAVI (Empirical Bayes Multi-omics Contrastive Analysis via Variational Inference), that leverages cross-modality relationships to impute RNA profiles for perturbations measured only by OPS, effectively generating Perturb-Seq data *in silico*. Moreover, EB-MoCAVI denoises and enhances sparse Perturb-Seq measurements with only a few cells per perturbation by leveraging matched imaging data that is more densely profiled.

PerturbPair demonstrated high overall concordance across modalities, but also fine perturbation-specific differences between RNA- and imaging-based analyses. OPS demonstrated superior sensitivity for detecting weak perturbation effects, likely due to its larger cell sampling capacity and targeted marker selection, whereas the rich phenotypic features captured in Perturb-Seq provided greater interpretability. Using EB-MoCAVI, we expanded the RNA compendium to 1,596 gene perturbations by imputing profiles for 1,082 perturbations identified by PerturbView, and validated these imputations in a secondary Perturb-Seq screen of 120 previously unseen perturbations. Using both measured and imputed profiles, we constructed a comprehensive regulatory map of macrophage activation, identifying novel regulators of key pathways and demonstrating how multi-modal integration can make large-scale perturbation studies more accessible and informative. PerturbPair represents a significant advance toward building comprehensive, multi-modal atlases of cellular perturbation responses.

## RESULTS

### PerturbPair enables parallel-modality perturbation profiling in primary macrophages

To systematically map how genetic perturbations affect both molecular and morphological responses of cells, we developed PerturbPair—a flexible framework that combines Perturb-Seq (or any multi-omics assay that allows for profiling of perturbations, here Perturb-CITE-seq^5,6^), and optical pooled screening (OPS) in the same biological system (**Figure 1A**). This approach, where each modality is performed in parallel, allows us to directly compare RNA and imaging phenotypes at scale, revealing which cellular processes are better captured by each modality and identifying perturbations that would be missed by either approach alone.

**Figure 1.**
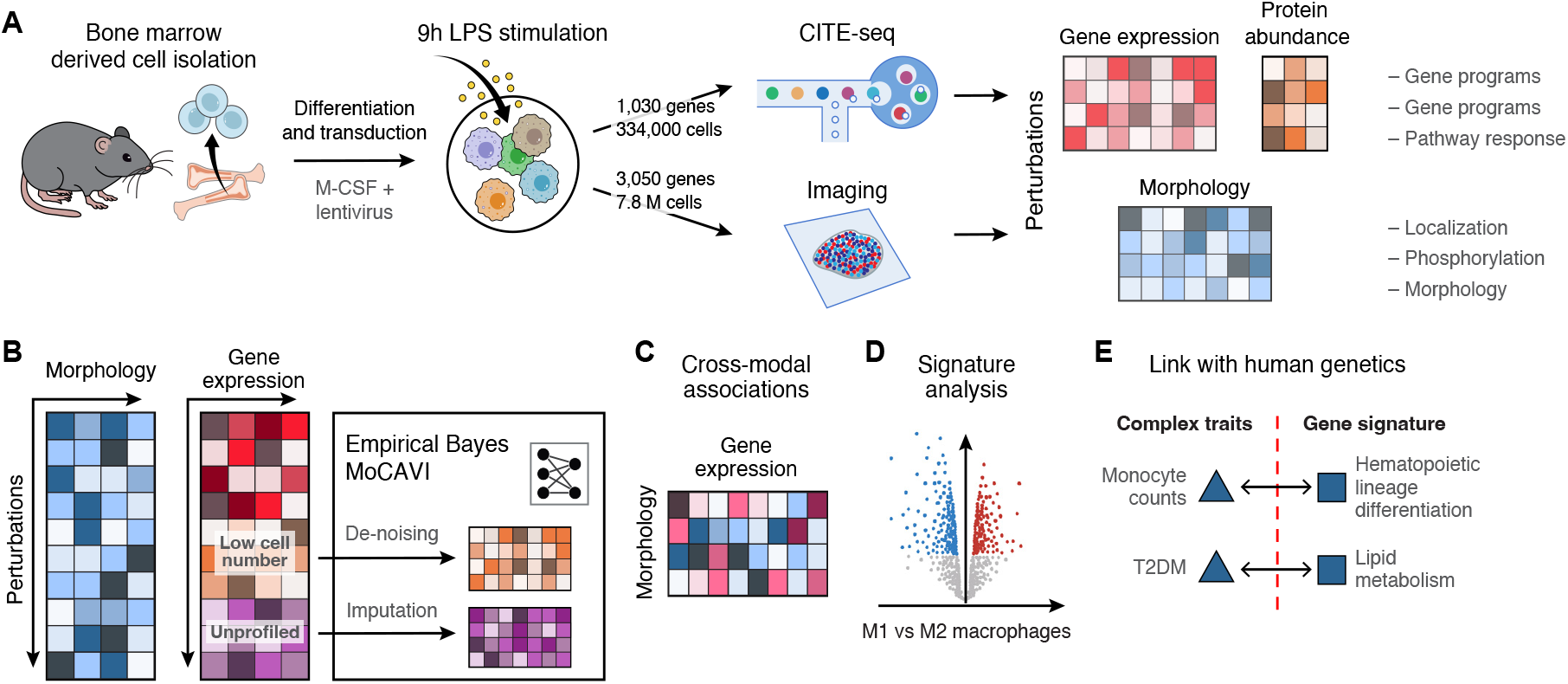
PerturbPair. (A) Experimental workflow. (B) Imputation of missing perturbations and denoising low cell number perturbations with EB-MoCAVI (Figure 4). (C-E) Additional computational capabilities from PerturbPair. (C) Relationships between imaging and RNA features. (D) In silico screening using gene expression signatures available in the literature, with applications to the characterization of regulators of macrophage states. (E) De novo identification of gene signatures for complex traits and disease after integration with rare-variant burden test loss-of-function data from UK Biobank (Figure 6).

We designed a CRISPR knockout (KO) perturbation library relevant to the response of primary BMDMs to LPS. For PerturbView, we used a 13,042 sgRNA library targeting 3,050 genes (4 guides per gene), spanning regulators of inflammatory responses^23^, phagocytosis^24^, and cellular morphology^13^, along with 412 negative controls targeting olfactory receptor genes and 40 negative controls that are non-targeting for the mouse genome. For Perturb-Seq, we used a subset of 4,114 sgRNAs targeting 1,030 of the 3,050 genes, with 300 negative controls targeting olfactory receptor genes and 40 non-targeting controls (**Table S1**). Both libraries were synthesized and cloned into a CROP-seq-based PerturbView backbone vector^14^, enabling dual readout.

To conduct the screens, we isolated bone marrow cells from Cas9 transgenic mice and differentiated them into BMDMs with M-CSF. On day 2, we lentivirally transduced them with each of the two libraries at a low multiplicity of infection (MOI = 0.2). After seven additional days of selection in puromycin and M-CSF, we stimulated mature BMDMs with lipopolysaccharide (LPS). Following nine hours of stimulation, one set of cells was harvested for Perturb-Seq (Perturb-CITE-seq)^25^, and another set was fixed with paraformaldehyde for OPS (PerturbView). We chose the 9-hour timepoint to capture both the transient nature of the inflammatory response and early protein changes^26–29^. We collected comprehensive dual-modality data to enable extensive computational analysis (**Figure 1**): 334,633 scRNA-seq profiles from Perturb-Seq (307 and 74 cells per gene and guide on average, respectively), and 7.8 million cells (2,576 and 638 cells per gene and guide on average, respectively) from PerturbView. Our Perturb-Seq analysis is specifically focused on cell profiles for 2,119 guides (547 genes) with significant effects (**Supplementary Note 1; Figure S1A-C**) and all controls, and the RNA measurements of 10,000 highly variable responding genes (RNA).

### Perturb-Seq recovered known and novel regulators of inflammatory signaling pathways

Primary BMDMs exhibit inherent expression heterogeneity that can confound the identification of perturbation-specific responses. For example, cell cycle states can vary naturally, in the absence of genetic perturbation, but also be affected by perturbations. Other cell state variation may arise only after genetic perturbation. Distinguishing these sources of variation is a topic of active research^30–34^. Here, we employed MoCAVI^35^, a variational inference framework for contrastive analysis that can operate across modalities (**Figure 2A, Methods**) and disentangle single-cell profiles into a background embedding that contains variations existing in non-perturbed BMDM populations, and a salient embedding that contains effects that are unique to perturbed cells (**Figure 2A**).

**Figure 2:**
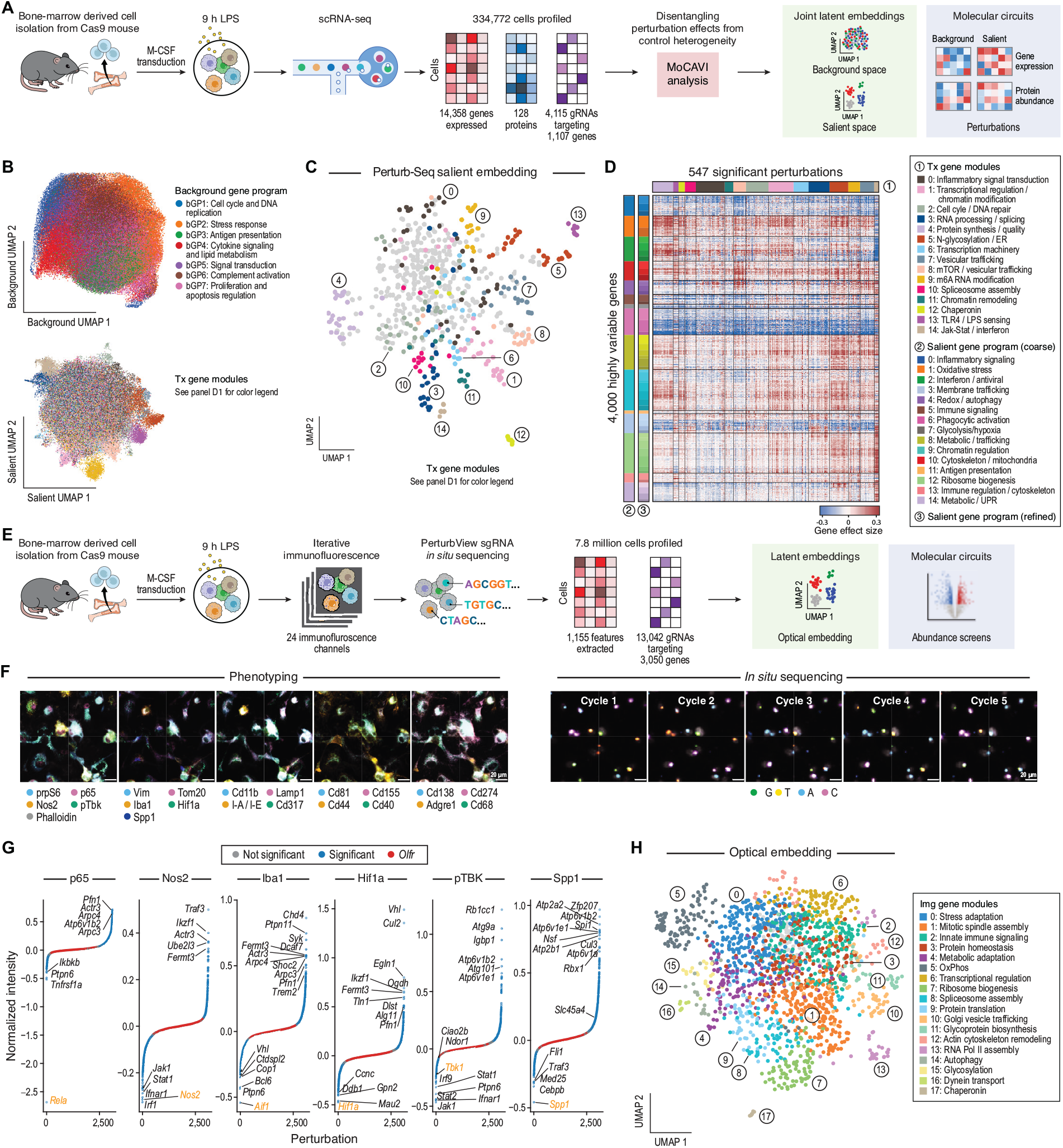
Perturb-Seq and PerturbView dissect gene networks involved in macrophage response to LPS. (A) Perturb-Seq analysis. (B) MoCAVI separates background and salient spaces. Uniform Manifold Approximation and Projection (UMAP) representation of Perturb-Seq profiles (dots) in MoCAVI’s background (top) and salient (bottom) spaces, colored, respectively, by the highest-scoring background gene expression program (top) or by the perturbation modules (bottom; legend in panel D). (C) Perturbation modules. UMAP representation of the pseudobulk profile of each gene perturbation (dot) in MoCAVI’s salient latent space, colored by gene module (legend in panel D). (D) Regulation matrix in the salient space. Regulatory effect sizes (color bar) in the salient latent space for each gene perturbation (columns) on the expression of each of the 4,000 most impacted genes (rows). Genes are clustered into coarse (color bar 2, far right) and fine (color bar 3, right) salient gene programs (GPs), and perturbations into co-functional modules (color bar 1, top). (E) PerturbView analysis. (F) PerturbView representative images. Representative images of iterative immunofluorescence for phenotyping (left) and in situ sequencing for genotyping (right). Scale bar, 20 μm. (G-H) Perturbation characterization by imaging phenotypes. G, Impact (normalized intensity, y axis) of each perturbation (x axis, ranked in increasing impact order) on either median (nuclear p65, nuclear Nos2, cellular Iba1, nuclear Hif1a) or maximum (cellular pTBK, cellular Spp1) protein levels. Highly impactful perturbations are highlighted. Orange: perturbation of gene encoding the measured protein. Blue/red: significant/non-significant perturbations. H, UMAP embedding of pseudo-bulk optical feature profiles (Methods) of each perturbation (gene level, dots), colored by OPS perturbation modules. See also related Figure S2.

MoCAVI identified seven background gene programs (bGP), with distinct functional features annotated by Hotspot analysis^36^ (**Figure 2B left, S1D-F, Table S2**). For example, bGP1 was enriched for cell cycle and proliferation genes (including Cdk1, Ccnb1, Ccnb2, Plk1, Mcm2-10, Cenpa, Cenpe, Cenph, Cenpf, and DNA repair components Rad51, Rad51b, Brca1, Brca2), and was most prominently induced in the Cdkn1a (p21) knockout (**Figure S1G**). This increased proliferative activity is consistent with p21’s role as a critical cell cycle checkpoint protein. This illustrates how pre-existing variation (e.g., cell cycle) can also be impacted by perturbation. Further interpretation of the background gene programs is provided in Supplementary Note 2 (**Figure S1H**).

MoCAVI’s salient space consisted of 15 modules of perturbations with similar functional effects impacting 4,000 genes partitioned into 15 coarse-grained gene programs (GPs), and 55 finer sub-programs (**Figure 2C-D, S1I-L, Table S3-4, Methods**). Guide pairs targeting the same gene or targeting two genes in the same complex or process (from CORUM or STRING^37,38^) had higher cosine similarity in the salient space than guides targeting random pairs of genes or guide pairs targeting the olfactory receptors, suggesting consistent effects for the targeting guides (**Figure S1I,J**). Module 13 captured positive regulators of TLR4/LPS signaling (Tlr4, Myd88, Irak4) whose perturbation strongly inhibited the LPS response, as expected (**Figure S1K**). Module 5 was enriched for N-glycosylation/ER genes (Alg1, Alg2, Ddost, Rpn1, Rpn2) whose perturbation dampened immediate-early NFκB responses (Tnf, Il1b, Il6, Nfkbia) but did not affect an ISG signature (Ifit1, Mx1, Gbp family) (**Figure S1K**). This aligns with the requirement for receptor glycosylation: N-glycosylation of Tlr4 and Md-2 enables LPS recognition and receptor assembly, phenocopying MyD88 disruption, yet preserving interferon routes^39,40^. Additional interpretations of the salient gene modules are in Supplementary Note 3 (**Figure S1K**). Relating the perturbation modules to the gene programs revealed distinct regulatory logic for each module (**Figure 2D**). For example, Module 14 (JAK-STAT pathway: Ifnar1, Irf9, Jak1, Stat1, Stat2) emerged as a master regulator with opposing effects: strong reduction of an interferon-stimulated gene program (GP2), and strong activation of the biosynthetic programs “ribosomal proteins and translation machinery” (GP12) and glycolysis (GP7). This reflects the trade-off between antiviral responses and cellular biosynthesis, where type I interferon signaling normally suppresses translation through JAK-mediated PKR activation to redirect resources toward antiviral defense^41,42^. Additional interpretations of the salient gene programs, as well as the surface protein programs are in Supplementary Note 3 (**Figure S1L**). Overall, the regulatory relationships demonstrate that macrophage responses involve coordinated regulation across multiple functional axes, with specific modules controlling distinct aspects of inflammatory and metabolic programs through separate biological mechanisms.

### Morphological profiling by PerturbView uncovered the regulatory architecture of LPS response

We conducted optical pooled screening with PerturbView, using 23 marker stains across five rounds of immunofluorescence, each round followed by fluorophore bleaching with the IBEX method (**Figure 2E**), and finally in situ sequencing (**Methods, Figure 2F**, right). Our panel included diverse morphological markers (DAPI, Phalloidin/actin, Vimentin, TOMM20) and key macrophage biology proteins: p65/RelA (NFκB), Nos2 (canonical M1 marker), Iba1/Aif1 (pan-myeloid), Spp1/osteopontin (immunomodulatory), and Hif1a (metabolic regulation) (**Figure 2F, left, Table S5**). After assigning sgRNAs and extracting features, we removed morphological outliers and guides represented by <150 cells, retaining 7.9 million cells out of the original 10.0 million cells detected (2,576 and 638 cells per gene and guide on average, respectively) (**Methods, Figure S2A-C**).

The screen was well-supported by internal controls and established biology. The 17 markers for which our library contains sgRNAs against the gene encoding them or their known regulators were depleted under those perturbations (empirical p-value; p < 0.05, **Figure S2D**, right), providing a positive control. For example, guides targeting p65 substantially decreased nuclear p65 intensity compared to guides targeting olfactory receptors (**Figure S2D, left**), and, the guides targeting the genes encoding each of p65, Nos2, Iba1, Hif1a, pTBK and Spp1 all ranked amongst the highest for perturbing the corresponding staining features (**Figure 2G**). Many of the regulators affecting each protein feature were consistent with known macrophage biology, as well as revealing new connections. For example, 711 regulators significantly impacted nuclear p65 intensity (empirical p-value; FDR < 5%), including Rela (p65) (strongest hit), and Tnfrsf1a, the TNF receptor (second strongest hit), consistent with the importance of TNF signaling in amplifying NFκB activity upon LPS stimulation, as well as key negative regulators, including several canonical signaling pathway members (e.g., Ptpn6, Ikbkb and Chuk) and transcriptional regulators (e.g., Nelfa and Dmap1), and positive regulators enriched for cytoskeletal and actin-associated proteins (Arpc3, Arpc4, Actr3, Pfn1). Additional biological interpretation for other stainings is in Supplementary Note 4 (**Figure 2G, S2E**).

A perturbation-specific embedding generated by principal component analysis with 1,155 extracted imaging features and 5,368 guides (1,596 genes) with significant and consistent effects (Methods) revealed a comprehensive regulatory architecture with multiple interconnected biological processes governing macrophage responses to LPS stimulation (**Table S6**). In the embedding space, cosine similarity between guide replicates targeting the same gene was significantly higher than between guides targeting olfactory receptor genes or random pairs of genes (empirical p-value; p < 0.01; **Figure S2F**). Additionally, cosine similarity between guides targeting genes in the same protein complex (from STRING or CORUM) was significantly higher than between those targeting random gene pairs (empirical p-value; p < 0.01; **Figure S2F**). Clustering the embedded perturbation profiles into 18 perturbation modules, showed that they contained coherent regulators impacting key processes (**Figure 2H, S2G**). For example, module 15 contains Jak/Stat pathway members, together with genes involved in m6A RNA methylation, dolichol-phosphate-mannose (DPM) synthesis and GPI-anchor biosynthesis. m6A RNA methylation was highlighted as a Nos2 regulator in recent screens^23^. Although DPM is known to donate mannose for diverse processes (including N-glycosylation), our results refine its mechanism of action: (i) the association of N-glycosylation genes with TLR/NFκB pathway rather than Jak/Stat pathway in our Perturb-Seq analysis (**Figure S1K**), and (ii) DPM genes co-cluster with the GPI-anchor pathway in one perturbation module. Together, these data support a model in which DPM loss dampens JAK– STAT signaling via impaired GPI-anchor biosynthesis, likely via lipid-raft perturbations that disrupt IFNAR nanoscale organization and endocytic trafficking. In another example, Slc45a4—a recently-described polyamine transporter implicated in pain perception in sensory neurons^43^—clustered with protein-folding genes in perturbation module 3 (e.g., Wdr81) (**Figure 2H**), which all increased Spp1 intensity (**Figure 2G, S2H**). This suggests that Slc45a4 influences Spp1 post-translational modification and secretory phenotypes in macrophages.

Next, we asked whether perturbation-perturbation similarities are conserved when solely using the four channels of broad morphology (DAPI, phalloidin (actin), Tomm20 (mitochondria) and Vimentin (intermediate filaments; similar to CellPainting^44^) vs. the full panel of 23 channels (which includes protein stains) (**Figure 2H, S2I**). Interestingly, there was a very high correlation of the energy distances in embeddings of profiles based on the four vs. 23 channels (**Figure S2J**; Spearman’s *ρ* = 0.89; p < 0.01). Additionally, embeddings from the reduced set of channels performed similarly on the CORUM/ STRING benchmark (**Figure S2K**). One notable difference was Rela perturbation whose effects were more noticeable by the full panel, as expected for p65 staining.

### Relationships between perturbations by OPS and Perturb-Seq are highly concordant

To compare the RNA and imaging modalities, we generated an RNA-specific embedding, as enabled by MoCAVI (**Methods**). Globally, the relationship between overall perturbation profiles was highly concordant between OPS and Perturb-Seq, when comparing the pairwise cosine similarities between perturbations for the 514 perturbations that had a significant effect in both modalities (**Figure 3A, S3A**). The perturbations grouped across the two modalities into modules enriched for genes involved in core cellular processes including protein complex assembly, transcriptional regulation, RNA processing, cytoskeletal dynamics, autophagy, and immune signaling pathways critical for macrophage activation.

**Figure 3.**
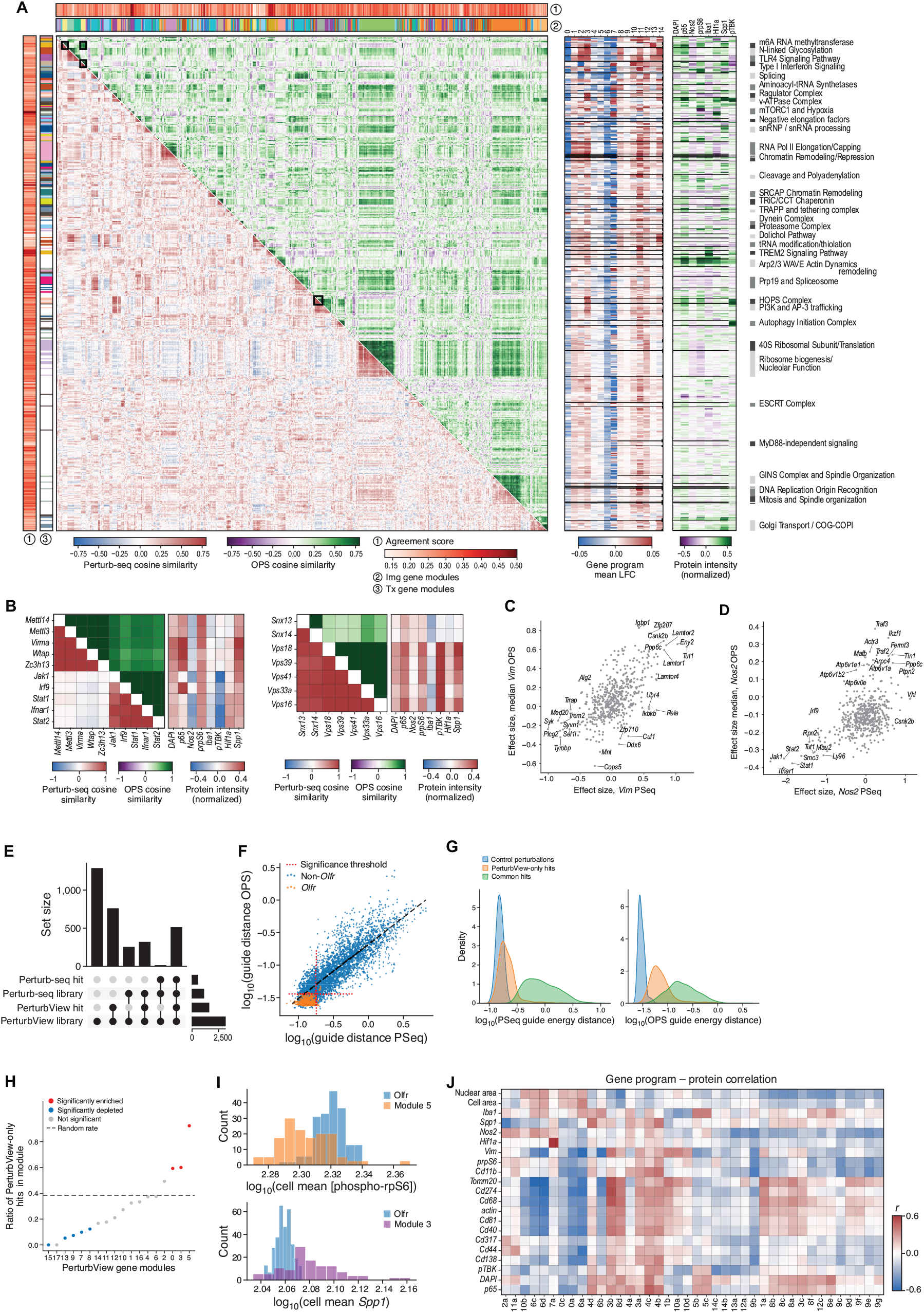
Joint analysis shows high concordance of regulatory architecture by Perturb-seq and PerturbView. (A-B) Similarity and distinction in co-functional regulators by Perturb-Seq and PerturbView. A. Cosine similarities between pseudobulk profiles of each gene perturbation (rows, columns) calculated separately by Perturb-Seq salient space embeddings (bottom triangle, red/blue color bar) or by PerturbView embeddings (upper triangle, green/purple color bar) (left matrix) and effect of each perturbation (rows) on gene programs (middle matrix, columns) or select optical features (right matrix, columns). Genetic perturbations are ordered by hierarchical clustering of joint sequencing and imaging phenotype vectors. Right-most and top-most red/white color bars: Agreement scores (Jaccard index of the 50-nearest neighbors overlap between each modality). B. Zoomed-in version (black box in A) with cosine similarities and protein staining intensity. (C-D) Relation between RNA and imaging features. C, Pseudobulk effect size of each perturbation (dot) on mRNA expression (x-axis, measured by Perturb-Seq) and protein staining levels (y-axis, measured by PerturbView) for vim/Vimentin (C) and Nos2/Nos2 (D). (E-G) Extent of agreement between screens. E, Top: Number (y axis) of unique and shared genes and hits in Perturb-Seq and PerturbView. Right: number of hits and genes in each screen overall. F, Perturbation strength (energy distance, x and y axes) for each guide present in both libraries (dots) based on either Perturb-Seq (x axis) or PerturbView (y axis) embeddings. G, Distribution of energy distances for guides present in both libraries based on Perturb-Seq (left) or PerturbView (right) data for shared hits (green), PerturbView-only hits (orange) and control guides (blue). (H-I) Perturbation modules enriched for PerturbView. Proportion of PerturbView-only hits (y axis) in each PerturbView perturbation module (x axis, ranked by increasing ratio). Dashed line: proportion of PerturbView-only hits across all considered perturbations (expected ratio under uniform assignment of perturbations to clusters). Enrichment and significance are assessed with a two-tailed hypergeometric test (p < 0.05). I, Distribution of mean intensity features (x axis) for marker proteins impacted by perturbation modules 5 (orange, top) and 3 (purple, bottom) that are significantly enriched PerturbView-only hits. Blue: Control guides targeting olfactory receptors. (J) Interpretation of protein imaging features with gene programs. Pearson correlation coefficient (color bar) between pseudobulk perturbation effects on protein staining intensity (rows, cellular median unless noted otherwise) and differential expression of gene programs estimated by MoCAVI (columns). Gene programs with at least one absolute correlation > 0.4 or at least three absolute correlations > 0.25 were kept for visualization. Maximum intensities were used for Spp1 and phospho-Tbk1. See also related Figure S3.

While the overall relationships between perturbations are highly concordant, at a finer level, there are differences based on RNA and imaging profiles. For example, JAK-STAT pathway components (Jak1/Irf9/Stat1/Stat2/Ifnar1) were highly correlated in one module in both modalities. However, Zfp207 and Tut1 clustered with this group only based on RNA (Perturb-Seq) profiles (**Figure 3B, left**), whereas m6A modification genes (Virma, Wtap, Zc3h13, Mettl3, Mettl14) only grouped with the JAK-STAT module based on imaging (PerturbView data). Indeed, while JAK-STAT perturbations strongly suppressed interferon-stimulated gene programs, m6A modification perturbations showed minimal impact on this program, suggesting that these pathways only converge post-transcriptionally (**Figure S3B**). In another example, endolysosomal pathway components showed distinct clustering patterns in each modality (**Figure 3B**, right). While perturbations to both HOPS complex members (Vps18/ Vps39/Vps41/Vps33a/Vps16) and to sorting nexins (Snx13/Snx14) were highly correlated in Perturb-Seq, the inter-group (HOPS-nexins) correlations in PerturbView were notably weaker. Analysis of interpretable imaging markers revealed functionally relevant distinctions: perturbations in Vps18/Vps39/Vps41/Vps33a/Vps16 strongly induced pTBK, HIF1a, and Spp1 accumulation, consistent with their essential role in global lysosomal fusion and the resulting blockade in protein clearance. In contrast, the Snx13/Snx14 perturbation group showed no distinct changes in the level of these proteins.

### Concordant and discordant perturbation effects on RNA and protein levels

To investigate the relationship between protein-level changes detected by OPS and RNA-level changes measured by Perturb-Seq, we compared perturbation effects for those proteins where we had both imaging channels and corresponding RNA data.

For some genes, there was high concordance between the effect of perturbations on their RNA and proteins, whereas for others, the agreement was low. For example, Vimentin RNA and protein responded to perturbations in a highly concordant way: perturbations that increased vimentin mRNA expression also increased protein abundance, and vice versa, e.g., mTOR pathway components (Tsc1, Tsc2, Lamtor1, Lamtor2, Rraga) (**Figure 3C**). Nos2 had moderate concordance: while the overall correlation was weaker than Vimentin, specific sets of perturbations showed concordant effects on both mRNA and protein (**Figure 3D**). The loss of JAK-STAT pathway components (Jak1, Stat1, Stat2, Ifnar1) led to concordant downregulation of Nos2 RNA and protein, reflecting the known dependence of Nos2 induction on interferon signaling pathways^23^, while the loss of TF Ikzf1 and signaling adaptors Fermt3 and Tln1 increased both RNA and protein levels, consistent with their roles in promoting inflammatory gene expression^45–47^. Conversely, the effect on Actin B (ActB) RNA and total protein level (phalloidin) was negatively correlated: perturbations that decreased median Actb RNA decreased total protein levels (**Figure S3C**). This may reflect homeostatic feedback mechanisms to maintain actin protein levels despite transcriptional perturbations, or to compensate for a change in protein degradation by increased or decreased transcription. For example, perturbation of Cul3, a component of E3 ubiquitin ligase complexes, which likely regulates actin protein degradation^48–50^, leads to increased Actb protein (e.g., through reduced degradation), but decreased RNA (**Figure S3C**). A similar pattern under perturbation of different vacuolar ATPases (Atp6v1a, Atp6v1b2, Atp6v1e1) may reflect the decoupling of transcription and proteostasis under lysosomal stress (**Figure S3C**). Finally, Hif1a showed a modest correlation (**Figure S3D**). Perturbation of Vhl, Cul2, and Egln1 caused larger increases in Hif1a protein levels, consistent with their known role in its posttranslational regulation: Vhl and Egln1 (PHD) mediate hydroxylation-dependent recognition of HIF1a^51^, while Cul2 drives the subsequent ubiquitination as the core of the ligase machinery^52^. This highlights how OPS can capture regulatory mechanisms that operate beyond transcriptional and RNA control. Overall, concordance between OPS and Perturb-Seq depends on the primary regulatory mechanism governing each gene perturbation, with structural proteins showing tight coupling and metabolic sensors exhibiting more complex post-translational control.

### OPS is more sensitive to detecting weaker perturbation effects due to higher cell numbers and specific stains

Despite the overall concordance, substantially more perturbations were “hits” (by statistical significance on the energy distance, **Methods**) in OPS than Perturb-Seq (**Figure 3E**). Specifically, of 1,030 genes perturbed in both experiments (hereafter, “Perturb-Seq library”), 514 were significant in both modalities (“common hits”), 321 only in PerturbView (“PerturbView-only hits”), and 16 only in Perturb-Seq (“Perturb-Seq-only hits”) (**Figure 3E**). This finding was unexpected given the high concordance in energy distance statistics at the guide level in the two assays (Spearman’s *ρ* = 0.85; **Figure 3F**) and in fitness, as reflected by cell proportions per perturbation (**Figure S3E**). However, PerturbView-only hits had low energy distance (in both assays) relative to other perturbations (**Figure 3G**), suggesting they elicit weaker effects in both readouts. Such weaker effects may require a larger number of cells for statistical significance, as available in PerturbView (**Figure S3E**).

Importantly, five of 18 PerturbView modules (#15, 13, 9, 7, 8), all related to some aspect of the core transcription, splicing, or translational machinery, were significantly depleted for PerturbView-only hits, and three were enriched (#5, 3, 0) (p < 0.05; hypergeometric test) (**Figure 3H**). The three modules enriched for PerturbView-only hits included perturbations whose effect can be sensitively detected by a post-transcriptional, cell biological readout. For example, Module 5 had the highest proportion of PerturbView-only hits, elicited a distinctive morphological phenotype, with depleted phospho-rpS6 (**Figure 3I**, top), and was enriched for genes encoding mitochondrial proteins (e.g., Mrpl54 and Ndufb8) and mTOR signaling components (e.g., Mtor, Rheb, and Rptor). We hypothesized that phospho-rpS6 staining, reflecting a post-transcriptional event in a key regulator of mitochondrial function that directly reflects mTOR activity, was particularly sensitive in detecting the impact of these perturbations (**Figure S3F**). In another example, Module 3 contained protein quality control and trafficking regulators (e.g., Wdr81) which elicited another distinct morphological phenotype. Perturbations in these genes disrupted the secretory pathway, causing intracellular accumulation of normally secreted proteins like Spp1 (osteopontin) (**Figure 3I**, bottom, **S2H**). This led to dramatic post-transcriptional changes of trapped proteins accumulating within cells, visible by immunofluorescence.

Together, these results highlight that the superior sensitivity of OPS for detecting hits with weaker effects arises from a combination of its inherent scalability—providing an order of magnitude more cells per perturbation—and its ability to capture complementary biological information through morphological features and targeted molecular markers that reflect primarily post-transcriptional events that may not have percolated to a strong RNA expression response.

### Describing imaging features from molecular programs

We next leveraged the shared perturbations to relate changes in imaging features to molecular changes in RNA-based gene programs. To this end, we calculated correlations between log-fold changes of fine-grained gene expression sub-programs and interpretable imaging features (median intensity of 19 markers unless otherwise noted) across the shared perturbations (**Figure 3J**).

This analysis mapped specific molecular axes (inflammation, proteostasis, metabolism, cytoskeletal remodeling) onto image-based phenotypes (**Figure 3J**). For example, the level of GP6 (6a, 6c, 6d; phagocytic activation) and GP0 (0a, 0c; inflammatory signal transduction) was negatively correlated with the levels of many inflammatory proteins (p65, Cd274, Cd44, Cd40, Cd68). GP6 was enriched for Fc gamma receptors and complement system genes (Fcgr1-4, C3, C1qa-c, Cfb, Cfp, C5ar1), which are often linked to tissue-repair functions^53,54^, whereas GP0 included negative feedback regulators induced by acute inflammation (Nfkbia, Socs3, Dusp1, Dusp2, Zfp36, Il10). In contrast, inflammatory GP3b (membrane trafficking) and GP4 (4b, 4c; proteostasis/autophagy) positively correlated with those pro-inflammatory markers, and encompassed processes that likely support inflammatory functions in antibacterial defense. Hif1a was strongly correlated with GP7a (glycolysis), consistent with Hif1a’s role as a master regulator of the metabolic switch to glycolysis^55,56^.

Spp1 protein levels were reciprocally regulated against inflammatory and antigen-presenting, Th2-polarizing programs, providing mechanistic context for its established role as a biomarker of tissue remodeling and fibrosis^57,58^. Specifically, Spp1 protein levels were positively correlated with expression levels of GP4d, identifying a specialized secretory gene program reflecting active vesicular packaging. GP4d spans the components of a secretory cascade including genes for production of tissue-remodeling factors (Osm, Igfbp7) and the requisite ER folding and post-translational modification enzymes (Edem2, Tgm2, Sumf2) and downstream vesicular transport machinery (Pld1, Rabgef1, Rapgef1, Ralgps1). Spp1 was also uniquely negatively correlated with the acute inflammatory activation program GP2a, and GP11a, a program of professional antigen-presenting cells (APCs) genes, characterized by high MHC class II expression (H2-Eb1, H2-Aa) and the production of Th2-recruiting chemokines and immunomodulators (Ccl17, Ccl22, Il4i1).

GPs 5b, 8e, 12c, and 12d were positively correlated with nuclear p65 abundance and were significantly enriched for p65 targets^59,60^ (**Figure 3J, S3G**). They represent the canonical effector functions of the active NFκB response including inflammatory output (GP5b: Tnf, Il12b, Ccl3), the homeostatic anti-apoptotic arm (GP8e: Kras, Stat5a, Raf1, Notch2), and the metabolic and biosynthetic requirements of activation via ribosome biogenesis and RNA processing (GP12c, 12d: Nol9, Wdr12, Mdn1, Ddx27, Wdr36, Utp18, Polr1f). Conversely, GPs 1a, 9 (9c, 9d, 9e, 9g), and 12 (12e, 12g) were enriched for p65 targets but decoupled from nuclear p65 abundance, characterizing the negative feedback and signal termination phase, involving signaling inhibitors (GP1a: Tnfaip3, Cyld, Ptpn2, Otud7b, Nfkbiz, Cited2, Cd200r4, Cflar, Nr1d2, Txnip), ubiquitin-mediated protein turnover machinery (GP9c, 9d, 9e, 9g: Birc2, Xiap, Itch, Cblb, Fbxw11, Cop1, Ube3a, Ubr5, Herc2, Herc4, Usp1, Usp3, Usp9x, Usp47, Senp1, Nae1, Cand1, Uba6Zc3h12c (Regnase-3)), and nucleolar biosynthesis with mitochondrial genes (GP12e, 12g: Polr1a, Ncl, Ddx21, Nop2, Gar1, Nop56, Nop58, Tomm20, Timm10, Timm9, Cox10m Mrpl12, Mrps31).

Finally, GPs 3 and 4 correlated strongly with nuclear p65 but were not enriched for its transcriptional targets (**Figure S3G**), suggesting they are coupled through an upstream trigger of p65 accumulation. GP3 defines a lysosomal-autophagic stress program enriched for lysosomal and trafficking genes (Tfeb, Npc1, Lyst, Gaa, Gba, Naglu, Tpp1, Atp13a2) and autophagy machinery (Optn, Map1lc3a, Sqstm1, Atg7, Plekhm1). GP4 constitutes an oxidative stress and immune-sensory signaling gene signature comprised of innate receptors and sensors (Ikbkb, Marco, Clec4d, Clec4n, Ly75), redox-defense genes (Txnrd1, Srxn1, Gclm, Prdx1, Cat, Mgst1, Slc7a11, Chac1, Sesn2), and inflammatory/metabolic effectors (Acod1, Osm, Cxcl1, Cxcl2, Cxcl3, Mmp12). These upstream programs were depleted of p65 binding sites, in contrast to their robust enrichment in negative feedback programs, likely reflecting a regulatory architecture that restricts positive feedback to ensure the transient nature of stress signaling^61^.

### EB-MoCAVI enables cross-modality imputation and denoising

Given the concordant yet complementary nature of RNA and imaging data revealed by our dual-modality screens, we next developed a computational framework that could leverage the relationship between modalities to address two key challenges: (1) imputing RNA responses for perturbations measured only by imaging, and (2) denoising RNA measurements when only a few cells are available per perturbation, by incorporating information from matched imaging data. To the best of our knowledge, this specific instance of perturbation denoising is a new task in the field of perturbation analysis. To this end, we developed EB-MoCAVI (Empirical Bayes Multi-omic Contrastive Analysis via Variational Inference), a machine learning framework that extends MoCAVI’s contrastive analysis approach with empirical priors derived from optical pooled screening data.

EB-MoCAVI takes as input a pair of datasets: genomics perturbation data (e.g., gene expression x_n_ ^g^ and guide RNA assignments t_n_ for cell n) from a subset of perturbations, and imaging perturbation data (in the form of pseudobulk perturbation embeddings e_r_) from a larger set of perturbations that includes both measured and unmeasured conditions in the genomics dataset (**Figure 4A**). EB-MoCAVI also supports additional modalities for the genomics dataset such as single-cell protein abundance or chromatin accessibility (**Figure 4A, left box and 4B, top left box**). This particular instance relies on CITE-seq measurements and also models protein levels y_n_ ^p^. The output consists of predicted RNA responses (or other ‘omics modalities) for unseen perturbations, denoised estimates of ‘omics profiles for perturbations with limited cell sampling, and latent space representations that capture both background cellular heterogeneity and perturbation-specific effects.

**Figure 4:**
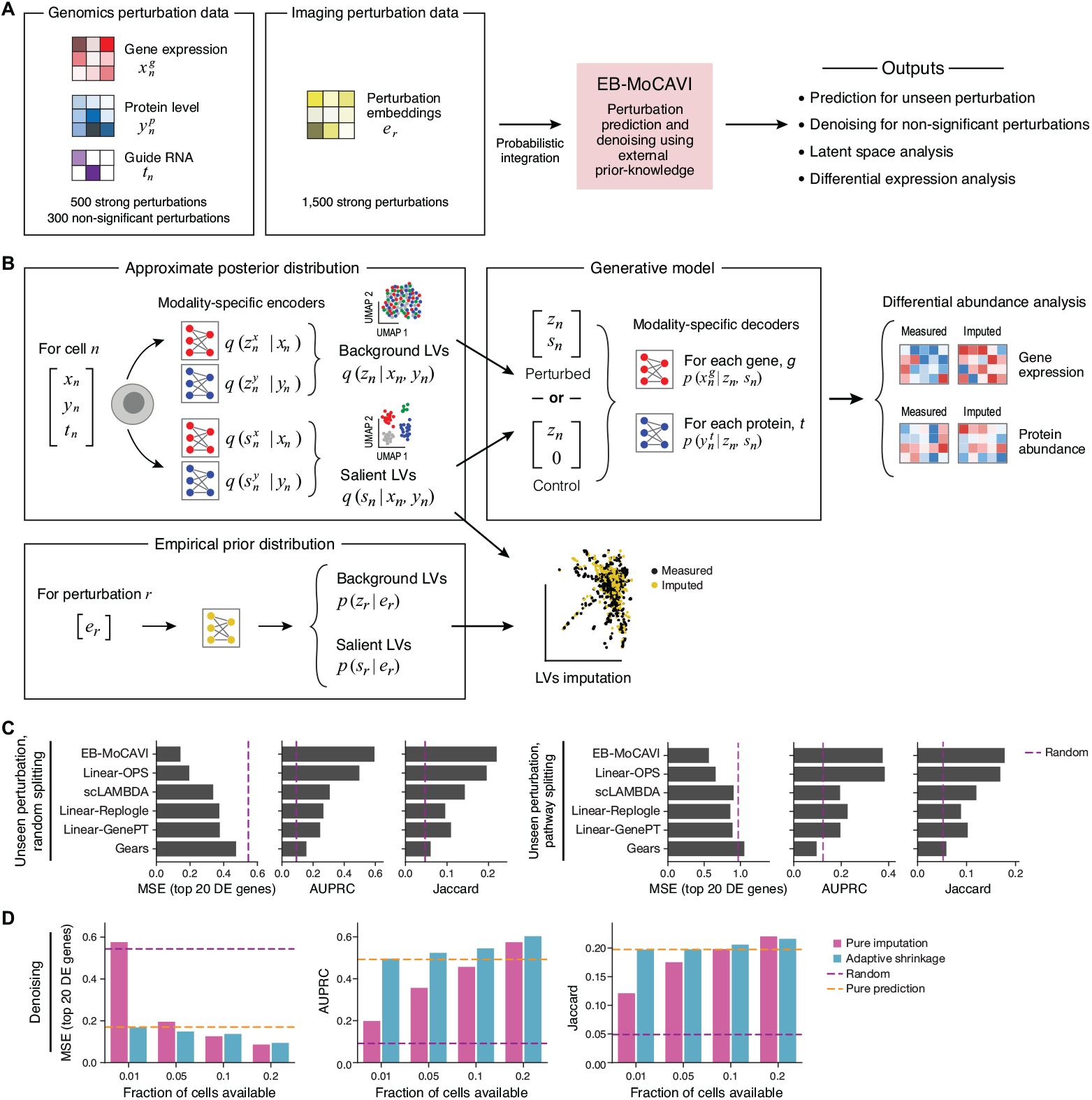
EB-MoCAVI framework for cross-modality imputation and denoising. (A) EB-MoCAVI concept. PerturbPair experiments yield two heterogeneous data sources. Genetic screens with single-cell readout (leftmost, e.g., Perturb-Seq) generate one or more sets of ‘omics profiles, in the form of count matrices, which for each cell n and gene g (or protein p, etc.) include the RNA expression level x_n_ ^g^ and the protein abundance y_n_ ^p^, along with the label t_n_ of the genetic perturbation in the cell (i.e., a guide RNA). OPS with imaging readout (second from left) generates a perturbation embedding, e_r_, for each perturbation r, based on imaging features. EB-MoCAVI (center, pink box, and panel B) takes these as input and infers the parameters of generative models for ‘omics profiles under perturbations, as well as latent variables from the data. It then uses these values to perform downstream analyses such as prediction of unseen perturbations, denoising as well as differential expression. (B) EB-MoCAVI generative model and inference approach. Data from each ‘omic modality is passed through two separate modality-specific encoder networks (top left box). The first set of modality-specific encoder networks encodes the parameters of the approximate posterior for the background latent variables z_n_, for the biological variation in the control population. The second set of encoder networks encodes the parameters of the approximate posterior for the salient latent variable s_n_, for the biological variation that is most affected by the perturbations. Next (top middle box), modality-specific decoder networks map the latent variables [z_n_, s_n_] to the parameters of a count distribution for every feature captured by the modality (e.g., gene, region, or protein). For any cell in the baseline environment (t_n_ = 0), the salient latent variable s_n_ is deterministically set to s_n_ = 0 before feeding into the decoder networks. The prior for latent variables [z_n_, s_n_] is defined via a prior network that takes as input the embedding e_r_ (bottom left box) . After training, the decoder networks may be used to characterize the effect of perturbations in both spaces, and the prior network may be used for imputation and denoising purposes (Methods). (C-D) EB-MoCAVI’s performance for gene expression predictions (imputation) or denoising. (C) Mean squared error for top 20 genes (MSE), Area under the precision-recall curve (AUPRC), and Jaccard similarity (x axes) for each method’s (y axis) predicted profiles vs. measured ones for perturbations that were heldout either randomly (left, random splitting) or for an entire held-out cluster (pathway splitting, Methods). Results are reported as the average across five distinct random seeds. (D) MSE, AUPRC, and Jaccard similarity (y axis) between fully measured profiles of held-out perturbations and either EB-MoCAVI’s denoising of measurements using a small fraction of cells (“adaptive shrinkage”, blue) or using only measurements of that fraction (“pure imputation”, pink), for increasing fractions (x axis). Orange dashed line: Denoising using only the prior (“pure prediction”). Purple dashed line: Performance of a “scrambled” predictor where values are sampled at random from the training data.

To model genomics data, EB-MoCAVI employs a variational autoencoder framework^62^ that disentangles cellular responses into background and salient latent variables^31^. For each cell n, the background latent variable z_n_ captures natural cell heterogeneity present even in control conditions (e.g., cell cycle in dividing cells), while the salient latent variable s_n_ encodes perturbation-specific effects (as in **Figure 2**). Similarly to previous work, the RNA level of each gene g follows a negative binomial distribution determined by a neural network decoder that maps the latent variables to RNA frequencies^62^. The key innovation of EB-MoCAVI lies in the specification of an empirical prior^63^: rather than using uninformative priors, both z_n_ and s_n_ are drawn from distributions parameterized by neural networks that take as input perturbation embeddings e_r_ from optical pooled screening (**Figure 4A, second box from left, Figure 4B, bottom left box**). This design enables the model to leverage rich morphological information to inform the predictions of ‘omics (e.g., RNA) profiles.

To impute responses for entirely unseen perturbations, EB-MoCAVI uses the empirical prior networks trained on imaging embeddings to generate latent variable distributions for perturbations not present in the genomics training data (**Figure 4B**). For denoising perturbations with limited cell counts, the model employs an adaptive weighting scheme that interpolates between prior-based predictions (from OPS embeddings) and posterior estimates (from measured Perturb-Seq data), with the weight determined by sample size (**Methods**). This approach enables a principled quantification of uncertainty and a smooth transition from prediction-dominated to data-dominated inference as the number of cells profiled by Perturb-Seq increases.

We benchmarked EB-MoCAVI’s perturbation effect imputation against several baseline methods (**Figure 4C**), including linear models using different perturbation embeddings (our OPS data, genome-wide Perturb-Seq data in K562 cells^64^, and GenePT language model embeddings^65^), as well as GEARS^66^, which is based on a graph neural network built from gene ontology pathways. We evaluated performance using three metrics: mean squared error on the top 20 most differentially expressed genes (MSE Top-20), area under the precision-recall curve for identifying differentially expressed genes (AUPRC), and k-nearest neighbors Jaccard similarity for preserving perturbation relationships (**Methods**). We either heldout perturbations randomly (at the gene level; “random splitting”), or, as a more challenging case, held out entire Perturb-Seq modules of co-functional perturbations (“pathway splitting”) (**Methods**). Across all evaluations, EB-MoCAVI consistently outperformed all baseline methods (**Figure 4C**), followed by a linear model using our measured OPS data, which also highlights the strength of the OPS data. Notably, the improvement was also significant when using pathway-informed splits, where all the perturbations from a given pathway are held-out in training to test generalization to functionally distinct perturbations. The results were robust across many random seeds, additional similar metrics (median absolute error instead of MSE, top 50 instead of top 20 genes), and did not require any hyperparameter optimization of EB-MoCAVI. This demonstrates the model’s ability to robustly capture meaningful biological relationships within the system.

For the denoising task, we systematically subsampled cells from randomly selected test perturbations to simulate scenarios with limited data availability, comparing three strategies within the EB-MoCAVI framework: pure prediction (using only OPS-based priors), pure imputation (using only available Perturb-Seq profiles), and adaptive shrinkage (combining both sources of information). The shrinkage approach consistently had the best performance across different levels of data sparsity, with the greatest improvements observed when very few cells were available (1-5% of original cell counts) (**Figure 4D**). Taken together, imaging data can serve as an effective regularizer for RNA profile inference, particularly valuable for large-scale perturbation experiments where comprehensive single-cell profiling may be cost-prohibitive.

### Imputation from OPS yielded a regulatory landscape of macrophage activation integrating imaging and RNA phenotypes

We used EB-MoCAVI’s imputation framework to predict gene expression profiles for the 1,082 gene perturbations, including 761 perturbations assayed only by PerturbView and 321 perturbations assayed in both PerturbView and Perturb-Seq but not significant in Perturb-Seq. We then merged these predictions with measured Perturb-Seq profiles for 514 gene perturbations to yield a joint dataset of 1,596 gene perturbations with either measured or imputed RNA phenotypes, which we embedded and visualized with PHATE^67^ (**Figure 5A**).

**Figure 5.**
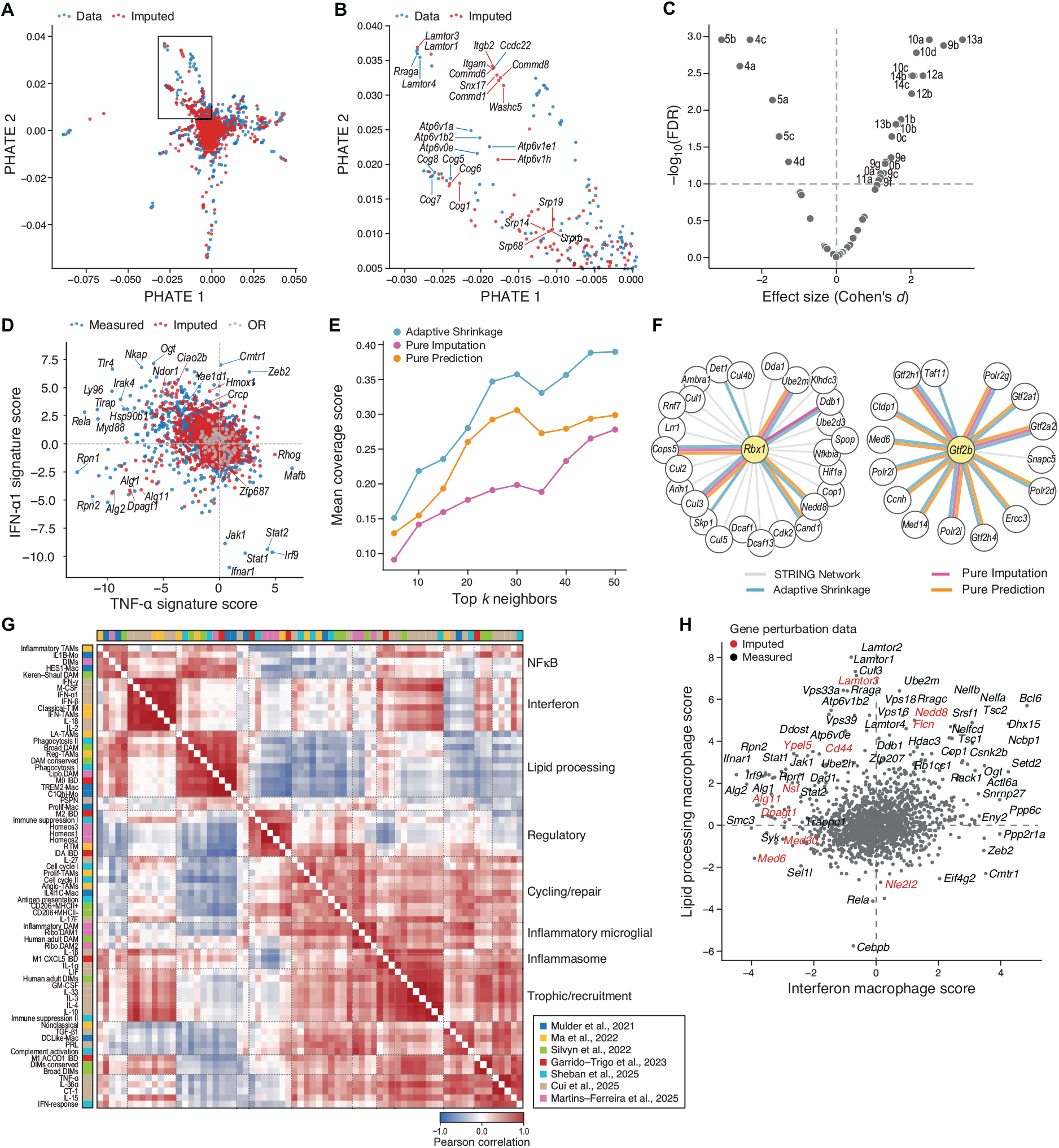
MoCAVI imputed and denoised Perturb-Seq profiles highlight regulator functions from a virtual or virtually-enhanced screen. (A-B) Integrated measured and imputed Perturb-Seq. A, PHATE embedding of measured (blue) and MoCAVI-imputed (red) pseudobulk Perturb-Seq profiles (dots). Dashed box: zoomed segment in (B). B, Zoomed-in of the dashed region in (A) with highlighted perturbation names. (C) Distinct features of a cluster of imputed perturbation profiles of SRP components. Effect size (Cohen’s D, x axis) and significance (-log_10_(FDR)) of mean log fold change (LFC) in program scores (dots) for the perturbations in the SRP cluster vs. their 28 neighbors. (D) Regulation of TNF-α and IFN-α1 signaling axis by measured and imputed regulators. Expression scores for the TNF-α (x axis) and IFN-α1 (y axis) programs for each measured (blue) and imputed (red) perturbation profile as well control (olfactory receptor targeting, gray). (E-F) Quality of denoised perturbation embeddings. E, Mean coverage score (y axis, proportion of top-k nearest neighbors in embedding space that are in the same protein complex as the query perturbation) for different values of k (x axis) for 34 PerturbView-only hits with <90 cells in Perturb-Seq, denoised by adaptive shrinkage (blue), pure prediction (orange), or pure imputation (red). F, Real protein-protein interactions (all edges, from STRING) for Rbx1 (left) and Gtf2b (right), detected as nearest neighbors by adaptive shrinkage (blue), pure prediction (orange), or pure imputation (red). (G-H) Prediction of regulators of key macrophage states. G, Pairwise Pearson’s correlation (color bar) between curated gene signatures (rows, columns; hierarchically clustered) based on their scores across measured and imputed perturbation profiles. Color bars: source studies. H, Mean signature scores for interferon macrophage (x axis) and lipid-processing macrophage (y axis) signatures in each measured (black) or imputed (red) perturbation profile. See also related Figure S4.

The imputed perturbation profiles clustered in a biologically coherent manner. Specifically, imputed profiles grouped with measured profiles from genes belonging to the same established complexes and pathways. For example, an imputed Lamtor3 perturbation profile localized alongside measured profiles for Lamtor1, Lamtor4, and Rraga perturbations, as expected for components of the lysosomal Ragulator–Rag axis that scaffolds mTORC1 activation (**Figure 5B**). Likewise, the imputed Atp6v1h perturbation profile was positioned among measured profiles for perturbations in other V-ATPase subunits. The imputed perturbations in members of the commander complex (Commd1, Commd6, and Commd8) clustered with measured profiles for Snx17, Washc5 (a WASH complex subunit), and leukocyte integrins Itgb2 and Itgam. Because Commander cooperates with SNX17 and WASH to recycle transmembrane cargos— including integrins—from endosomes to the plasma membrane^68^, co-localization with measured perturbation of Ccdc22 (a core CCC component linking COMMD proteins to Commander) further supports the biological integrity of this module in our embedding (**Figure 5B**).

In particular, we observed a distinct grouping consisting only of imputed perturbation profiles of Srp14, Srp19, Srp68, and Srprb (the β subunit of the SRP receptor) (**Figure 5B**). Because the signal recognition particle (SRP) directs nascent secretory and membrane proteins to the endoplasmic reticulum, defects in SRP components are known to trigger Regulation of Aberrant Protein Production (RAPP)—a co-translational surveillance pathway that selectively degrades mRNAs encoding secretory/vesicular clients to prevent accumulation of mistargeted proteins^69^. We therefore hypothesized that SRP perturbations would suppress specific expression programs as a result of RAPP activity and its downstream effects. To test this at neighborhood resolution, we identified 28 perturbations (19 measured and 9 imputed) in the same Leiden cluster as the SRP profiles and compared program-level mean log-fold changes between the SRP perturbation profiles and those neighboring perturbations (**Figure 5C**). GPs 4a, 4c, 4d (Redox / autophagy) and 5a, 5b, 5c (immune signaling) were significantly more downregulated by SRP perturbations (FDR < 0.05) than by the neighboring perturbations. These results suggest a failure to engage the ER stress and UPR responses, consistent with RAPP-mediated mRNA degradation eliminating the luminal inputs required to activate these sensors.

Next, we identified both measured and imputed perturbations that modulate the downstream TNF-α and IFN-α1 signaling axes induced by LPS by scoring gene expression against gene signatures derived from macrophages stimulated *in vivo* with these specific cytokines^70^ (**Methods**). While both pathways are pro-inflammatory, their expression levels were generally anti-correlated across perturbations (**Figure 5D**), consistent with the well-defined mutual antagonism between TNF-α and interferon pathways^71^. Perturbation of genes encoding JAK-STAT pathway components decreased the expression of the IFN-α1 signature while increasing TNF-α, highlighting a negative feedback loop where basal interferon signaling normally restrains TNF-α-driven inflammation. In contrast, several measured perturbations of genes encoding core NFκB pathway components (Tlr4, Rela, Myd88, Hsp90b1, Ly96, Tirap) decreased expression levels of the TNF-α signature but increased levels of the IFN-α1 signature. Based on several imputed profiles, perturbations of genes involved in N-glycosylation (Alg11, Dpagt1) attenuated both signatures, consistent with measured perturbation profiles of other N-glycosylation regulators (Alg1, Alg2, Rpn1, Rpn2) (**Figure 5D**). Among the most impactful hits upregulating the IFN-α1 signature were the imputed perturbation profiles for Hmox1, Ciao2b, and Ndor1, together with the measured perturbation profile for Yae1d1 (**Figure 5D**)—key components of the cytosolic iron–sulfur assembly (CIA) pathway that delivers Fe–S clusters to nucleocytosolic proteins from Hmox1-driven iron supply to Ndor1-mediated electron transfer and Ciao2b/Yae1d1-directed cluster delivery^72,73^. Their convergence suggests that compromising Fe–S maturation/redox transfer primes macrophages for interferon response. Mechanistically, defects in Fe–S biogenesis and redox homeostasis can increase reactive oxygen species (ROS), biasing cells toward heightened inflammatory signaling^74^. These findings nominate redox/Fe–S homeostasis as a tractable axis for tuning macrophage inflammatory state and illustrate how the expanded OPS–Perturb-Seq compendium uncovers biology beyond the directly assayed space.

We assessed EB-MoCAVI’s denoising, on the subset of 34 PerturbView-only hits which had only 23-90 cells per gene perturbation in Perturb-Seq, asking if it improves their biological interpretability compared to pure prediction (using only OPS) or pure imputation (using only Perturb-Seq data) approaches (**Methods**). We benchmarked the performance of each approach at embedding perturbation profiles in a manner consistent with known protein-protein interactions, by calculating the fraction of the top-k most similar perturbations that were also neighbors according to STRING and CORUM. Across all tested neighborhood sizes (k=5 to k=50), EB-MoCAVI’s adaptive shrinkage consistently outperformed both pure prediction and pure imputation, achieving mean coverage scores that were 30-60% higher than pure imputation (**Figure 5E**). For example, Gtf2b (40 cells) achieved coverage scores of 0.93 versus 0.27 for pure imputation, successfully identifying 14 of 15 known transcriptional machinery partners including RNA polymerase II subunits (Polr2d, Polr2g, Polr2i, Polr2l) and other general transcription factors (Gtf2a1, Gtf2a2, Gtf2h1) (**Figure 5F**, right). Similarly, Rbx1 (79 cells), a core component of cullin-RING ubiquitin ligases, showed improved recovery of known partners including Cul3, Skp1, Nedd8, and Cops5, relationships that were largely missed when relying solely on the limited measured Perturb-Seq data (**Figure 5F**, left). To further assess our denoising, we identified the top-5 unique neighbors by cosine similarity in the salient space and examined those neighbors that were detected only after denoising, but not after either pure prediction or pure imputation. For example, Copb1 (42 cells in Perturb-Seq) was a top-5 neighbor of Cog6 (measured in OPS only), consistent with the biochemical cooperativity of β-COP (Copb1) and the COG complex, including COG6, in Golgi trafficking^75,76^. In another example, Snrnp200 (34 cells in Perturb-Seq) was a top-5 neighbor of Ess2 (measured in OPS only), both established components of the spliceosomal C complex^77^. Thus, OPS data serve as a powerful regularizer for inference of RNA profiles, enabling biologically meaningful perturbation representations and associations even when Perturb-Seq data are very limited.

### Large-scale virtual Perturb-Seq nominates regulators of recurrent macrophage states

One of the strengths of Perturb-Seq vs. OPS is high phenotypic interpretability on a transcriptome scale, which is easier to integrate with prior knowledge such as in vivo cell atlases from patients or functional gene signatures. To quantify how gene signatures co-vary across perturbations, we scored 514 measured and 1,082 imputed Perturb-Seq profiles with a curated compendium of macrophage signatures from diverse experimental conditions, tissues, and disease states^70,78–83^, and then calculated pairwise correlations between the signatures based on their score profiles (**Figure 5G**). The impact of genetic perturbations organized the signatures into a few clusters, suggesting that macrophages adopt a limited set of recurrent states across contexts. A similar analysis using only profiles from BMDMs carrying sgRNAs targeting olfactory receptors showed much weaker patterns (**Figure S4**), indicating that the covariation revealed by Perturb-Seq reflects biologically meaningful impact of the perturbations, rather than only baseline BMDM variation.

One prominent cluster was enriched for interferon responses: signatures of macrophages from IFN-γ–, IFN-α–, and IFN-β–injected mice grouped tightly with interferon-primed tumor-associated macrophages (TAMs) (**Figure 5G**, “interferon”). Another consisted of signatures from TAMs, disease-associated microglia (DAMs), macrophages in inflammatory bowel disease, phagocytosis gene programs, and lipid-associated macrophages (LAMs) (**Figure 5G**, “lipid processing”). This suggests convergence on a phagocytic, lipid-remodeling state that appears in both tumors and damaged neuroinflammatory tissue.

We then used the scores to identify measured or imputed perturbations that can shift cells between these interferon or lipid-processing states (**Figure 5H**). As expected, disruption of JAK–STAT components and N-glycosylation genes decreased interferon-program scores. Conversely, perturbation of mTOR-pathway regulators or V-ATPase subunits (e.g., imputed Lamtor3 and measured Lamtor1, Lamtor2, Lamtor4, Rraga, Rragc, Atp6v1b2, Atp6v0e) increased lipid-remodeling scores, consistent with mTOR/V-ATPase control of lysosomal acidification and degradative capacity^84^. Notably, knockout of Zeb2, Cmtr1, or Eif4g2 (all measured) uniquely increased interferon scores, while suppressing lipid-remodeling scores. Zeb2 has recently been implicated as a master regulator of the inflammatory status of TAMs ^81^. Cmtr1 and Eif4g2 influence selective mRNA translation, highlighting translational control as a lever to restrain inflammation and to switch macrophages between states^85^.

### Regulator-program analysis revealed regulators and gene programs of human myeloid lineage traits

Perturb-Seq has recently emerged as a powerful approach for dissecting the biological mechanisms underlying human genetic associations^86–88^. Following recent approaches^86^, we integrated our measured and imputed Perturb-Seq profiles with rare-variant loss-of-function (LOF) burden test statistics to assess its utility for identifying trait-relevant gene programs and putative trait-specific upstream regulators (**Figure 6A**).

**Figure 6.**
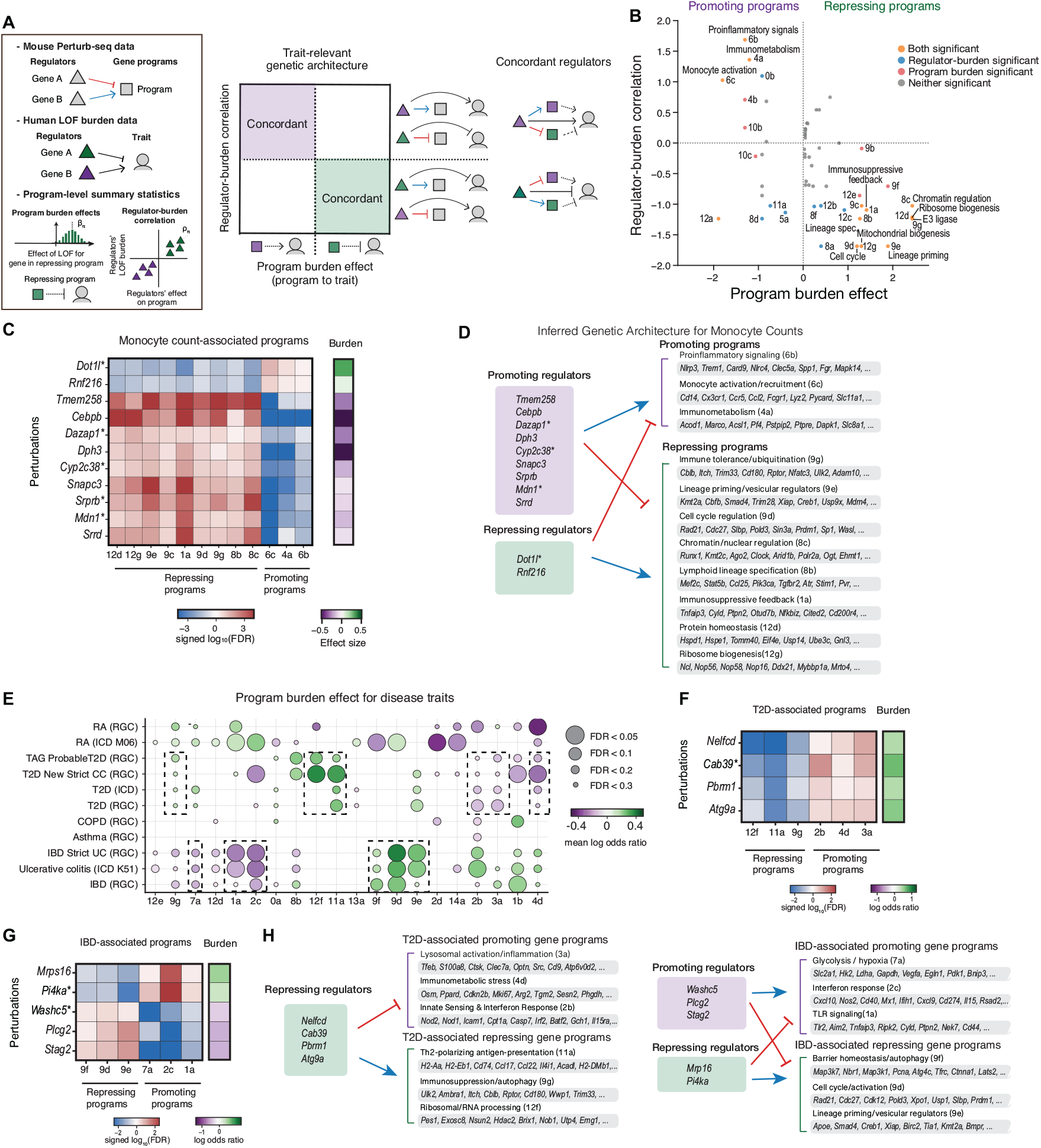
Co-analysis of Perturb-Seq with loss-of-function burden tests highlights causal regulators of human traits and diseases. (A) Analysis approach. Left: Perturb-seq data (top) are used to determine whether a gene (triangle) up-regulates (blue, solid pointy arrow) or down-regulates (red, solid blunt arrow) downstream expression programs (squares). LOF burden effects from human genetics data (middle) quantify whether a gene is trait-repressing (green triangle) or a trait-promoting (purple triangle). From these data, two program-level summary statistics are computed (bottom). Program burden effect γ_π_ quantifies whether a gene program promotes (purple square, dashed pointy arrow) or represses (green square, dashed blunt arrow) a trait. Regulator burden correlation tests whether positive (blue, solid pointy arrow) and negative (red, solid blunt arrow) regulators’ effects on a program match their LOF burden associations with the trait. Right: Regulators of multiple programs were nominated as concordant if their LOF burden association with the trait was directionally consistent with their effects on significant programs. (B-D) Regulators and programs associated with monocyte counts. (B) Program burden effects (signed log_10_(FDR), empirical p-value, x-axis) and regulator-burden correlation (signed log_10_(FDR), empirical p-value, y-axis) of each of 55 sub-programs (dots) in monocyte cell count trait from UKBB, colored by significant associations on program burden effects (red), regulatory-burden correlation (blue) or both (orange). (C) Effect of program-wide concordant regulators (rows) on mean LFC of gene programs (columns) (signed log_10_(FDR), empirical mean p-value of LFCs, left heat map) and on monocyte counts (log odds ratio, right). Asterisks: Imputed perturbation profile. (D) Schematics illustrating causal relationships between positive (blunt arrow) and negative (blunt arrow) regulators, gene programs and the trait (here, monocyte counts). (E) Gene programs associated with disease traits. Program burden effect (dot color; mean log odds ratio) and significance (dot size, FDR) of each program (columns) across 10 disease traits (rows). (F-H) Key regulators of T2D- and IBD-associated gene programs. (F,G) Effect (signed log_10_(FDR)) of each selected regulator (rows, left) on each gene program (columns, left) and on disease risk (log odds ratio, right) for T2D (F) and IBD (G). Asterisks: Imputed perturbation profile. (H) Schematic of key regulators of T2D (left) or IBD (right)-associated gene programs. See also related Figure S5.

Briefly, in this approach^86^ we take as input two relations: first, *γ*_g_ denotes the effect of gene *g*’s LOF on the trait through burden association (**Figure 6A, dashed pointy/blunt arrows; Methods**); second, perturbseq data provide estimates for β_g→π_ : the effect of knocking out gene *g* on the expression score of program *π in vitro* (positive/negative regulators by Perturb-Seq; **Figure 6A left, solid pointy/blunt arrows**), for each of the 56 gene sub-programs. Next, to identify trait-relevant programs, we compute two key summary statistics for every program *π* : a program burden effect *γ*_*π*_, and a regulator-burden correlation *ρ*_*π*_ (^86^; **Methods, Figure 6A left**). We define the **program burden effect** *γ*_*π*_ as the mean effect *γ*_*g*_ of the genes *g* that are members of program *π* on the trait, with significance tested by permutation. Positive values indicate that the program genes’ on average repress the trait (repressing programs, **Figure 6A green square and blunt dashed arrow**), and negative values indicate that the program genes on average promote the trait (promoting programs, **Figure 6A purple square and pointy dashed arrow**). We define, for each program and trait pair, the **regulator-burden correlation** *ρ*_*π*_ as the correlation between *β*_*r*→*π*_ (the effect of knocking out each regulator *r* of program *π* on the expression score for that program) and *γ*_r_ (the repressing/promoting effect of that same regulator on the trait through LOF burden association), using only regulators with significant effects in at least one of the analyses (i.e., FDR < 0.05 for either program regulation or LOF burden). As a proof of concept, we focused on the quantitative trait of monocyte counts from the UK Biobank (UKBB) cohort, given its relevance to our experimental model. We collected LOF rare-variant burden statistics^89^ and denoised them using GeneBayes^90^ (**Methods**). For comparison, we ran the same analysis pipeline on lymphocyte counts, eosinophil counts, and neutrophil counts.

Thirteen gene programs were significantly associated with monocyte counts in both program burden effects and regulator-burden correlation (**Figure 6B,C, Figure S5**) and eight of them were also associated with neutrophil counts (of 10), but few if any were associated with lymphocyte counts (2 regulator significant programs) or eosinophil counts (0 significant programs) (**Figure S5**). This aligns with the closer developmental origin of monocytes and neutrophils to granulocyte–monocyte progenitors (GMPs)^91^ vs. the common lymphoid progenitors (CLPs) for lymphocytes^92^ and the eosinophil-lineage– committed progenitors (EoPs) for eosinophils^93^. For both monocyte and neutrophil counts, most programs were concentrated along the anti-diagonal line, consistent with concordant program–regulator associations (i.e., loss of program genes and loss of their activators have effects in the same direction, whereas loss of repressors would have effects in the opposite direction), which supports the idea that these programs may be causally interpreted^94^ (**Figure 6B**).

In three cases, both the program burden effect and regulator-burden correlation were significant for promoting monocyte counts (**Figure 6B, top left quadrant, orange**). GP6b contains pro-inflammatory signaling genes (Il17ra, Plk3, Naaa, Fgr kinase, Pak1, Clec5a), including genes from the inflammasome pathway (Nlrc4 and Nlrp3) and Trem1 and Trem3, myeloid receptors that amplifies inflammatory signaling, reflecting acute and amplified inflammation. GP6c contains cell cycle regulators (Ccnd3, Nek6, Incenp), monocyte markers and inflammatory genes (Cd14, Fcgr1, Ccl2, Ccl7, Irf7, Il1rn, Ccr5, Cx3cr1), involved in classically-activated monocytes possibly undergoing expansion. GP4a contains genes involved in metabolic adaptation for immune functions (Acsl4, Marco, Slc16a10, Slc8a1, Slc48a1), as well as Irg1 (Acod1), which has been highlighted as a key regulator of inflammation in chronic inflammatory disorders^95^.

In contrast, repressing programs associated with reduced monocyte counts, many of which were also negatively associated with neutrophil counts, typically comprised of genes related to immunoregulatory functions and hematopoietic lineage regulation (**Figure 6B, bottom right quadrant**). For example, GP1a encompasses various negative feedback regulators of inflammation (Tnfaip3, Cyld, Ptpn2, Otud7b, Nfkbiz, Cited2, Cd200r4, Cflar, Nr1d2, Txnip) with immunosuppressive functions, opposite to the inflammatory functions enriched in promoting programs. GPs 8b and 8c, while both stem-cell associated, consist of genes that regulate distinct aspects of the progenitor state. GP8c genes maintain the epigenetic landscape of the hematopoietic stem cells (Runx1, Arid1b, Kmt2c, Clock), whereas GP8b genes drive lymphoid lineage specification (Mef2c, Stat5b, Ccl25), effectively biasing progenitors away from monocyte differentiation, consistent with the human genetic association. GP9g genes reinforce immune tolerance through ubiquitin-mediated feedback (Cblb, Itch, Trim33) and autophagy (Rptor, Ulk2, Ambra1). GP9e includes transcriptional and vesicular regulators, integrating progenitor maintenance factors (e.g., Kmt2a, Cbfb, Smad4, Elk4) and critical vesicular trafficking and metabolic genes (e.g., Eps15, Pikfyve, Apoe, Xdh), while GP9d consists specifically of DNA replication machinery and mitosis genes (Rad21, Cdc27, Pold3, Slbp). Finally, GP12g is enriched for ribosome biogenesis factors (Ncl, Nop56, Nop58, Ddx21), while GP12d features genes regulating protein homeostasis and mitochondrial entry (Hspd1, Tomm40, Usp14, Eif4e). These two programs are negatively correlated with monocyte counts but not neutrophil counts, highlighting the intense biosynthetic capacity of these precursor states.

To provide a genetic architecture of the trait, we next identified trait-specific putative upstream regulators. Because each trait-associated program was regulated by perturbations of many genes *in vitro*, we focused on 11 upstream regulators *r* whose rare-variant LOF associations with monocyte counts (*γ*_r_) were directionally concordant with their Perturb-Seq effects (*β*_*r*→*π*_) across all significant monocyteassociated programs *π*. Specifically, a regulator with *γ*_r_ > 0 was retained only if *β*_*r*→*π*_ > 0 for all monocyte-promoting programs (GP6c, 4a, and 6b) and *β*_*r*→*π*_ < 0 for all monocyte-repressing programs (GP12d, 12g, 9e, 9c, 1a, 9d, 9g, 8b, and 8c), whereas a regulator with *γ*_r_ < 0 was retained only if it showed the opposite pattern (**Figure 6C, D**). For example, LOF of Cebpb, a known regulator of monocyte development and maintenance^96–98^ is a strong suppressor of monocyte counts, and KO of Cebpb increased monocyte-repressing programs while reducing monocyte-promoting programs. Conversely, LOF in Dot1l (an imputed perturbation), the sole H3K79 histone methyltransferase, was associated with increased monocyte counts. Prior studies show that Dot1l sustains lipid biosynthesis programs and restrains inflammatory responses in macrophages and its genetic or pharmacologic inhibition shifts macrophages toward a hyper-inflammatory state^99^. Myeloid-specific Dot1l deletion in mice in vivo aggravated atherosclerotic inflammation and was accompanied by a higher proportion of circulating pro-inflammatory monocytes^99^. Consistent with these observations, early clinical experience with the DOT1L inhibitor pinometostat documented leukocytosis in 20% of patients and histological evidence of myeloid differentiation^100,101^. Thus, our framework’s prediction agrees with genetic, in vivo, and clinical perturbation data, with Dot1l loss reducing multiple monocyte-repressing programs (GP12d, 12g, 9e, 9c, 1a, 9d, 9g, 8b, and 8c) and increasing multiple monocyte-promoting programs (GP6c, 4a, and 6b), thereby supporting a causal chain from Dot1l to program to trait.

### Perturb-Seq ties human disease genetics to gene programs and regulators

Next, we assessed the relevance of our gene programs to several human diseases where macrophages have been implicated in disease etiology^102,103^: Type 2 diabetes, asthma, chronic obstructive pulmonary disease (COPD), rheumatoid arthritis, and inflammatory bowel disease (IBD), including ulcerative colitis. Because those diseases had low prevalence in the UK Biobank (from 0.9% for ulcerative colitis to nearly 11.1% for asthma), we collected rare-variant LOF burden scores^89^ and applied minor allele count filters to ensure sufficient statistical power for each disease (**Methods**). We focused on interpreting each disease trait’s program burden effect in each Perturb-Seq gene (sub)-program (**Figure 6E**, FDR<0.2).

Clustering by program burden effects grouped similar diseases together (**Figure 6E**). For example, IBD and ulcerative colitis formed a distinct cluster characterized by programs regulating antimicrobial defense and inflammatory signaling. GP2c, positively associated with disease risk, is defined by interferon-stimulated genes (ISGs; Mx1, Rsad2, Ifit family) and chemokines (Cxcl9, Cxcl10). Notably, this program contains Irgm1 and Sp140, established IBD susceptibility genes linked to autophagy and bacterial handling. GP1a, also a positive predictor, is enriched for negative feedback regulators of NFκB and cytokine signaling, including Tnfaip3 (A20), Ptpn2, Cyld, and Nfkbiz. Complementing these inflammatory drivers, GP7a represents a critical metabolic adaptation defined by aerobic glycolysis and HIF1α signaling (e.g., Hk2, Slc2a1, Vegfa). In contrast, GPs 9d, 9f, and 9e were negatively associated with IBD risk, highlighting a failure of specific homeostatic checkpoints. GP9d defines a post-transcriptional regulatory network for mRNA decay and silencing (Prdm1, Rc3h1, Cnot6), while GP9f represents a proteostasis and stress-response module driven by ubiquitin and autophagy regulators (Map3k7, Atg4c, Fbxo30). Furthermore, GP9e was enriched for tissue-survival and tolerance signaling, containing TGF-β pathway genes (Smad4, Bmpr2) and anti-apoptotic factors (Birc2). This pattern supports a model where susceptibility is driven by the unchecked activation of inflammatory and glycolytic programs, compounded by a concurrent failure of the post-transcriptional and proteostatic checkpoints necessary for immune resolution.

Type 2 diabetes (T2D) was positively associated with immunometabolic programs driving chronic cellular stress, GPs 2b, 3a, and 4d. GP2b encompasses genes encoding innate immune sensors (Nod1, Nod2) and apoptotic regulators (Casp7). GP3a defines a lysosomal stress module driven by Tfeb alongside lipid and iron handling mediators (Sptlc2, Ncoa4), and GP4d features Osm (Oncostatin M), a key cytokine driving tissue inflammation and insulin resistance^104,105^, alongside prostaglandin receptors (Ptger2, Ptgir) and oxidative stress regulators (Sesn2, Chac1). Together, these profiles suggest that persistent innate activation and maladaptive organelle responses to metabolic overload may promote macrophage-mediated T2D pathogenesis. In contrast, GPs 11a, 12f, and 9g were negatively associated with T2D disease risk, enriching for protective homeostatic functions. GP11a represents a Th2-polarizing, tolerogenic APC-like signature (H2-Eb1, Ccl17, Il4i1)^106^, whereas GP9g features vesicular trafficking and mTOR-regulated autophagy (Rptor, Ulk2, Ambra1) alongside E3 ubiquitin ligases (Itch, Cblb) implicated in immune tolerance^107,108^. GP12f comprises ribosome biogenesis and RNA processing genes (Utp4, Pes1, Exosc8, Nsun2), as well as genes for metallothioneins (Mt1, Mt2) that buffer oxidative stress. Thus, while IBD risk is driven in part by specific failures in pathways that limit inflammation in macrophages, T2D-associated gene programs in macrophages align with a broader immunometabolic dysfunction, defined by the impairment of the protein synthesis and autophagic machinery required to buffer inflammatory stress.

Rheumatoid arthritis (RA) risk was positively associated with programs orchestrating synovial inflammation and tissue remodeling. GP2d defines the core inflammatory engine of the joint, and comprises genes encoding the canonical M1 transcription factor Irf5 alongside genes for critical signaling kinases (Map3k8) and downstream effectors (Stat3, Casp8). GP14a consists of Unfolded Protein Response (UPR) genes, including the master regulator Xbp1 and ER chaperones (Hspa5, Calr), which may reflect the immense protein-folding capacity required by the hyperactive synovial macrophage^109^. In contrast, GP1a and 2c which were positively associated with IBD risk were negatively associated with RA risk, highlighting a tissue-specific divergence in macrophage regulation. While GP1a was enriched for anti-inflammatory genes (Tnfaip3), it also included critical survival and licensing factors (Cflar, Nfkbiz). These programs illustrate a pathogenic shift where the loss of intrinsic regulators permits the sustained inflammation and high secretory demand of the rheumatoid joint.

Using the same putative-regulator framework as above, we next examined program burden effects in T2D and IBD to identify regulators of the three disease-promoting and three disease-repressing gene programs in each disease, retaining those with concordant Perturb-Seq effects and significant LOF burden effects (FDR < 0.2; **Figure 6F–H**). For T2D, our analysis nominated four key regulators that govern metabolic adaptation and transcriptional regulation (**Figure 6F**). First, Cab39 (MO25) is the obligate scaffold protein required to stabilize the LKB1-STRAD complex and activate AMPK. While high-fat diet exposure dissociates Cab39 from LKB1, thereby uncoupling the cell’s energy sensor^110^, Cab39 overexpression restores AMPK signaling and suppresses NFκB-driven inflammation^111^. Next, Atg9a negatively regulates the cGAS-STING pathway by controlling STING translocation from the Golgi to endosomes^112^, such that Atg9a deficiency leads to hyper-responsive interferon signaling upon cytosolic DNA stimulation^112^. Third, Nelfcd is a transcriptional regulator that suppresses interferon-stimulated genes^113^, and its loss has been linked to increased efferocytosis and aberrant transcriptional response in response to flu and LPS^114–116^, suggesting that removing this molecular brake accelerates the clearance of apoptotic cells in the diabetic niche. Finally, Pbrm1 (BAF180) is a subunit of the PBAF chromatin remodeling complex. Although its specific function in macrophages has not been described, its established role in modulating tumor immunity^117,118^ and regulating ROS production in epithelial cells^119^ suggests that it may function as a buffer against metabolic and oxidative stress in macrophages in the diabetic niche as well.

Similarly, for IBD, we identified six key regulators essential for barrier defense and microbiome sensing (**Figure 6G**). Plcg2 (Phospholipase C Gamma 2) is a central signaling hub that transduces calcium signals for ITAM-coupled receptors, including Dectin-1 and TREM2^120,121^, and its LOF is associated with reduced risk of IBD, mirroring the protective effects observed with pharmacological inhibition of its upstream activator, BTK^122^. Stag2, a Cohesin complex subunit, helps organize the 3D genome to facilitate the rapid induction of inflammatory programs, a mechanism established in myeloid progenitors^123^. Pi4ka (Phosphatidylinositol 4-Kinase Alpha) is the enzyme for generating the phosphoinositide PI4P at the plasma membrane. Mutations of Pi4ka and its partner TTC7A are established as a cause of severe, early-onset IBD and immunodeficiency^124–126^.

Taken together, our analysis reveals that macrophage gene programs and their regulators, as defined by Perturb-Seq, display disease-specific patterns of rare-variant burden and functional association across a broad range of chronic inflammatory diseases. Using T2D and IBD as examples, we identified a sensible set of regulators of disease-associated gene programs (**Figure 6H**). These findings illustrate the value of interpretable cellular profiles (Perturb-Seq) for systematically linking gene program activity to human disease phenotypes, demonstrating a promising approach for the identification of critical regulatory nodes.

### A follow-up Perturb-Seq screen validates the accuracy of EB-MoCAVI’s predictions

Finally, we performed a targeted secondary Perturb-Seq screen profiling 42,265 cells across 288 gene perturbations including 117 perturbations selected with EB-MoCAVI-imputed profiles, 143 perturbations measured in the primary screen and 28 Olfr controls (**Figure 7A, S6A,B, Table S7**). For each target gene, we selected the single most potent guide based on the OPS screen. Notably, 88% of the 405 perturbations significantly impacted the transcriptome (**Figure S6C**). For the 143 perturbations with measured data in the primary screen, LFCs across 4,000 genes were highly reproducible across screens (median Spearman’s ρ=0.72; **Figure 7B, S6D**).

**Figure 7:**
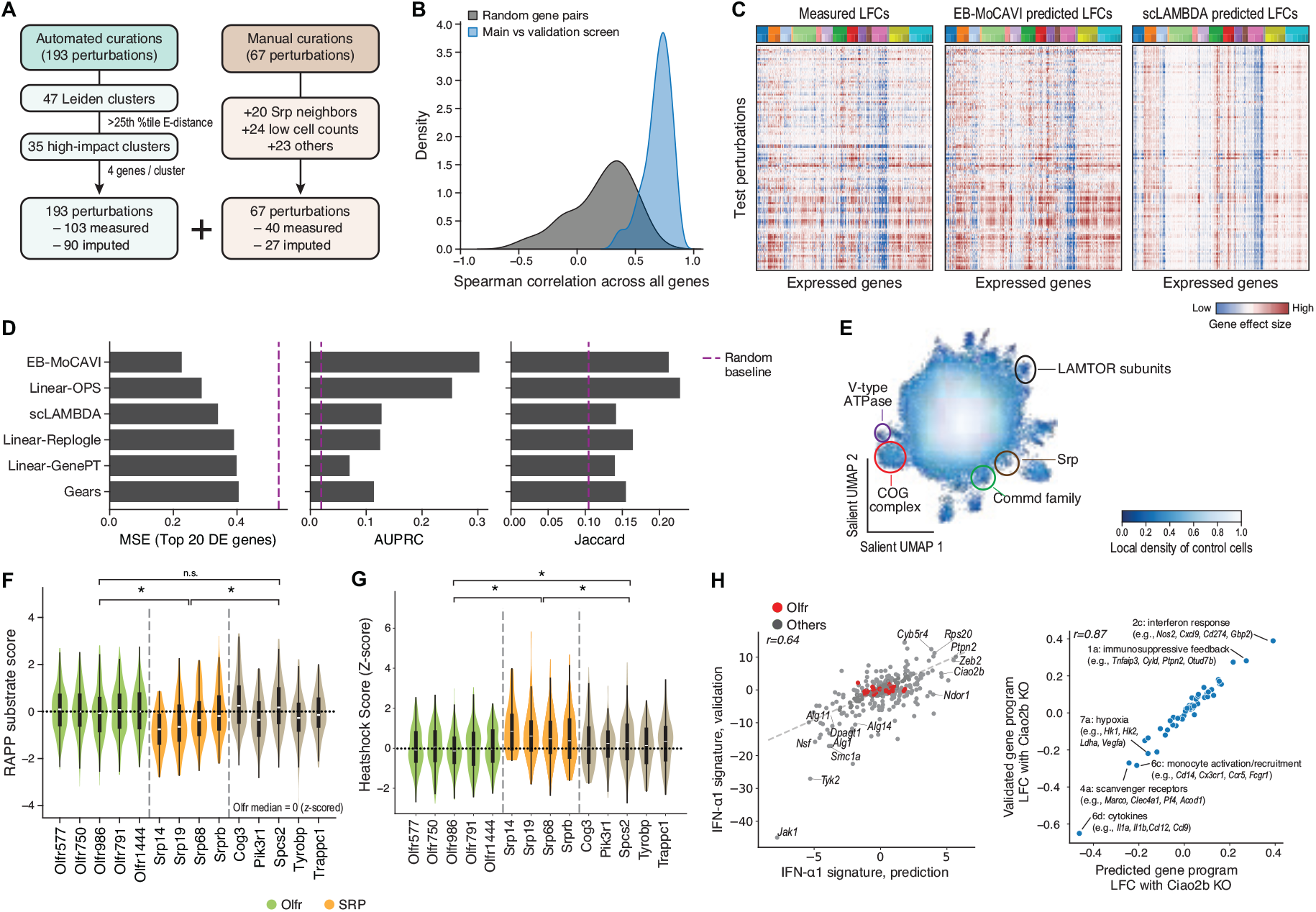
A secondary Perturb-Seq screen validates EB-MoCAVI’s imputations and denoising-based predictions. (A) Selection of perturbations for the secondary validation screen. (B) Screen reproducibility. Distribution of Spearman correlation coefficients (x axis) between 143 pseudobulk perturbation profiles measured in both the primary (main) screen and the secondary (validation) screen (blue) or between random pairs of pseudobulk perturbation profiles measured in the two screens (gray). (C-D) Validation of EB–MoCAVI’s imputation accuracy. C, Regulatory coefficients of the impact of 120 perturbations (present only in the secondary screen, rows) on 4,000 expressed genes (columns) based on measured secondary screen data (left), MoCAVI imputations of unmeasured profiles using the primary screen (middle), and scLAMBDA predictions from the primary screen (right, the best method that does not use OPS data). D, Mean square error for the top 20 genes (left), area under the precision-recall curve (AUPRC) (middle), and Jaccard similarity (right) (x axes) for each method’s (y axis) and 120 perturbations with predicted profiles from the primary screen vs. measured ones for the secondary screen. Results are reported as the average across five random seeds. (E-H) Validation of specific imputation-based predictions. E, UMAP of MoCAVI’s salient embedding of measured perturbation profiles (dots) from the validation screen, colored by density of control profiles. (Model trained on the validation data only). (F-G), Distribution of gene signature scores (y axis) for RAPP substrate genes (F) and heat shock response genes (MSigDB MM14951) (G) in cells carrying control (Olfr, green), SRP (gold), and neighboring (gray) perturbations. * P<0.05; n.s. P>0.05, empirical p-value. H, Ciao2b knockout upregulated type I interferon response. (left) IFN-α1 signature scores in pseudobulk perturbation profiles imputed from the primary screen (x axis) and measured in the secondary screen (y axis). Red: control (Olfr) cells. Upper left: Pearson’s r. (right) gene program-wise log-fold change of Ciao2b perturbation profiles imputed from the primary screen (x axis) and measured in the secondary screen (y axis). See also related Figure S6.

The secondary screen validated EB-MoCAVI’s imputation capabilities and the specific predictions drawn from imputed profiles. First, across all perturbations unseen in the primary screen, the imputed profiles closely recapitulated the observed RNA profiles (**Figure 7C**, median Spearman’s ρ = 0.48). This substantiates the performance of EB-MoCAVI’s imputation, which consistently outperformed scLAMBDA and other methods (**Figure 7C,D**). The expression profiles of the secondary screen also validated the functional clusters based on measured and imputed profiles from the primary screen (as described in **Figure 5B**), including LAMTOR subunits, V-type ATPase, COG complex, Commd complex and Srp family (**Figure 7E, S6E**). We further validated the specific local effects of SRP perturbations (as predicted in **Figure 5C**). To this end, we scored the measured profiles from the secondary screen against a curated set of RAPP substrates—defined as genes containing a signal peptide^127^ that are downregulated upon Srp54 knockdown^128^. To ensure specificity, we compared the SRP perturbation profiles against olfactory receptor controls and the non-SRP members of the same cluster in the primary screen (Cog3, Pik3r1, Spcs2, Tyrobp, Trappc1). We observed a significant and specific decrease in RAPP substrate expression solely in the SRP perturbations (**Figure 7F**). Furthermore, the loss of Srp genes led to a significant increase in heat-shock response compared to both control groups (**Figure 7G**)^129,130^; this upregulation of chaperones is consistent with the cytosolic accumulation of precursor proteins, a hallmark of SRP defects. Moreover, the empirical measurements from the secondary screen also largely validated our predicted IFN-α1 signature scores (Pearson’s r = 0.64) (**Figure 7H**). Consistent with our predictions, the imputed perturbations of N-glycosylation genes (Alg11, Alg14, Dpagt1) successfully captured attenuated inflammation. Similarly, the validation screen confirmed our prediction that iron-sulfur cluster loss drives interferon response; specifically, Ciao2b knockout resulted in high pathway activity and significant upregulation of inflammatory genes like Ifit3 and Nlrp3 (**Figure 7H, S6F**).

We also validated EB-MoCAVI’s denoising. To this end, we compared the imputed LFCs (derived solely from OPS data) and denoised LFCs against the secondary screen for 11 perturbations that yielded low cell counts in the primary screen but sufficient cell counts in the secondary screen (at least 20 cells). The remaining 13 perturbations intended to validate the denoising did not pass this threshold on the number of cells, and therefore could not be used for this evaluation. Denoising with adaptive shrinkage, on the sparse primary screen superiorly recovered the perturbation effects measured in the secondary screen (**Figure S6G**). We also confirmed the functional similarity of Copb1 to Cog5 and Cog6 (as predicted above) based on the strong correlation of gene program scores of denoised profiles from the primary screens and measured profiles from the secondary screen (**Figure S6H**). Specifically, the loss of each of Copb1, Cog5, and Cog6 upregulated ER stress (GP5a) and UPR activation (GP14a) programs while downregulating phagocytotic programs (GPs 6a, 6c, 6d, 10c), consistent with their established roles in retrograde trafficking machinery.

## DISCUSSION

PerturbPair advances our ability to map cellular perturbation responses by enabling direct comparison of ‘omics (e.g. RNA) and imaging (e.g. morphological) phenotypes within the same biological system. Aligned with previous work directly comparing imaging and sequencing data^20,131,132^, our dual-modality platform revealed both overlap and complementarity in the biological signal captured by the two readouts. Unlike previous work, we demonstrated for the first time that the concordance between those heterogeneous assays is high enough to rely on one of those modalities for cross-modal predictions. Looking forward, technological advances in single-cell multiomics may eventually enable capture of both transcriptomic and imaging information from the same individual cells, potentially providing even richer perturbation atlases while reducing experimental complexity. So far, existing studies are limited in the scale of perturbations or expressed genes that are profiled^21,133^. Moreover, the parallel nature of PerturbPair means that as new screening approaches arise, they can easily be included in the same framework.

Our dual-modality screen illustrated the utility of high-dimensional, multi-scale functional genomics to dissect the regulatory architecture of macrophage activation. Through this approach, we mapped how macrophages coordinate diverse biological processes and positioned specific regulators within a global hierarchy of functional potency and module connectivity. For example, by integrating protein level and localization (from imaging) and RNA levels (from scRNA-seq), we found that SPP1 abundance positively correlated with a secretory gene program but negatively correlated with inflammatory and antigen-presentation functions, highlighting a functional divergence between Spp1 protein expression and inflammatory gene programs. Furthermore, we dissected the regulation of p65, showing that nuclear p65 abundance marked a coordinated NFκB response in which oxidative stress and immune sensory programs reflected upstream activation, cytokine, metabolic, and ribosomal programs reflected downstream effector output, and distinct p65 target-enriched programs captured negative feedback and signal termination. Beyond individual signaling pathways, we defined recurrent multivariate cell states—such as an interferon-primed versus a lipid-processing phenotype—and identified specific regulators that shift cells between them. Notably, we identified translational regulators (Cmtr1, Eif4g2) and the transcription factor Zeb2 as key drivers of this transition. Our analysis also uncovered unexpected metabolic checkpoints, such as the cytosolic iron-sulfur assembly (CIA) component Ciao2b, which acts as a critical restraint on inflammatory tone by preventing hyper-activation of the interferon response. Ultimately, these findings demonstrated that scaled integration of perturbation signatures at the morphological and molecular level is essential for resolving the complex, multi-layered regulatory logic that dictates macrophage fate and function.

We introduced EB-MoCAVI, an empirical Bayesian framework^134^ that leverages optical pooled screening data for imputation of missing perturbation effects in single-cell genomics measurements. Perturbation prediction is known to be a difficult problem, for which performance highly depends on the available source of prior knowledge^135–137^. We demonstrated here for the first time that morphological information from the same biological system strongly improves performance for this task, outperforming all of the state-of-the-art methods we tested. This proof of concept has broader implications for the design of future multi-modal perturbation experiments, suggesting that OPS data can serve as an effective foundation for guiding more expensive single-cell genomics efforts. In this framework, a smaller Perturb-Seq screen can be coupled to an OPS screen which is 10-fold larger (or more), to generate a comprehensive view of the system. While our current analysis only focused on pseudo-bulked imaging data at the perturbation level for computational convenience, we anticipate that EB-MoCAVI could be improved with a full single-cell treatment of the imaging data as well, using frameworks we and others have recently developed to match distributions of perturbation data^138–142^.

The denoising capabilities of EB-MoCAVI address a persistent challenge in perturbation screening: the trade-off between perturbation coverage and statistical power per perturbation. Our adaptive shrinkage approach, which interpolates between prior-based predictions and single-cell data based on available sample size, achieved 30-60% improvements in biological relationship recovery for low-cell perturbations compared to RNA profiles alone. This advance suggests a shift toward designing perturbation experiments with broader coverage and lower per-perturbation cell counts ^2^, relying on computational methods to achieve adequate statistical power. However, realizing this potential will require development of improved uncertainty quantification methods that can provide calibrated confidence intervals for denoised estimates, enabling researchers to appropriately weight evidence from computationally enhanced versus directly measured perturbations^143^. Such advances could greatly democratize large-scale perturbation screening by reducing the cell numbers required per perturbation while maintaining biological interpretability. This should further allow a much faster exploration of both perturbation space and diverse biological systems, both of which are likely needed to develop generalizable “virtual cell” models for biology^1,22^.

The integration of our perturbation atlas with human genetics data demonstrates how cellular screening platforms can illuminate disease mechanisms at scale. Our systematic analysis revealed disease-specific gene program patterns—such as ER stress pathways in inflammatory bowel diseases and mitochondrial dysfunction programs in type 2 diabetes—that individual gene associations might miss. However, our current approach faces limitations when applied to binary disease traits, as the log-odds ratios from rare-variant burden tests exhibit higher noise levels than quantitative traits, reducing our ability to detect program-level associations. Future work could address this limitation by developing joint models that simultaneously analyze both perturbation and genetics data, potentially leveraging the cellular response patterns to improve genetic effect size estimates while using human genetic variation to validate perturbation-derived biological insights. Despite these limitations, the framework establishes a foundation for using perturbation atlases to systematically connect cellular mechanisms to human disease. As perturbation datasets expand across cell types and disease-relevant contexts, this approach promises to transform our understanding of how genetic variation manifests through cellular programs to influence human health, ultimately enabling more precise therapeutic targeting of disease-associated pathways.

## Author contributions

T.K. and R.L. and A.Re. conceived the study. T.K. and R.L. led the project, and wrote the manuscript with input from all authors. T.K., R.L., A.M.M., K.G.S., and A.S. developed the experimental design. A.M.M., P.C., and V.C. performed the BMDM isolation and contributed to the experimental design. T.K., A.M.M., and A.Ri. performed the Perturb-Seq experiments. T.K. performed the PerturbView experiments. R.L. led the theoretical formulation and developed the EB-MoCAVI framework, with input from T.K. and J.C.H. R.L. and T.K. performed the data analysis, with support from D.C.L. R.L. and T.K. conducted the human genetics analysis, with guidance from M.O. and J.K.P. K.G.S., A.S., and A.Re. jointly supervised the project, with additional guidance provided by O.R.R., L.G., and J.K.P.

## Acknowledgements and Disclosures of Funding

The authors thank David Richmond, Burkhard Hoeckendorf, Josh Kaminker, Joshua Gould, Jose Sergio Hleap Lozano and Bo Li for contributions to image processing; Yichen Gu, Russell Littman, and Sandra Melo-Carlos for helpful discussions on multimodal analyses; and Huisheng Zhu and Nikhil Milind for input on human genetics. We thank Jon Scherdin and Lindsay Liang for cloud computing and data storage support. The authors thank the NGS team in Genentech for next-generation sequencing processing support. The authors thank Leslie Gaffney for her help in figure making. Schematic figures were created with BioRender.com.

## Data and Code Availability

All raw and processed count data will be made available in the National Center for Biotechnology Information’s Gene Expression Omnibus upon publication of the manuscript. Raw imaging data are currently being deposited into a public repository upon publication. The implementation of EB-MoCAVI along with tutorials is publicly available on GitHub (https://github.com/Genentech/MoCAVI). EB-MoCAVI is implemented within the scvi-tools framework^146^.

## METHODS

### Animal experiments

All protocols involving animals were approved by Genentech’s Institutional Animal Care and Use Committee, in accordance with guidelines that adhere to and exceed state and national ethical regulations for animal care and use in research. All mice were maintained in a pathogen-free animal facility under standard animal room conditions (temperature of 21°C ± 1°C, humidity between 55 and 60% and 12-h light/12-h dark cycle).

### Cell culture

HEK 293T cells were all obtained from ATCC and were maintained in DMEM supplemented with 10% fetal bovine serum (FBS; vol/ vol), 100 U/ml penicillin, 100 μg/ml streptomycin and 1× GlutaMAX.

For BMDM isolation and culture, bone marrow was collected from tibias and femurs of 6- to 12-week-old male Cas9 transgenic mice ^147^. Following red blood cell lysis (ACK lysis buffer, Gibco, A1049201), 20 million cells were plated in BMDM medium (DMEM high glucose, 10% Tet-Negative heat-inactivated FBS, 1× GlutaMAX, 100 U/ml penicillin–streptomycin, and 100 ng/ml recombinant mouse macrophage colony-stimulating factor (Genentech media facility) at a density of 0.8 million cells per ml in 150-mm non-tissue culture-treated dishes (Corning). Cells were transduced on day 3 by adding lentiviral supernatants overnight in fresh BMDM medium. Following transduction, cells were split on day 3 and selected with 5 μg/ml puromycin from days 6 to 9. At day 12, cells were harvested and plated onto 6-well glass-bottom plates (CellViz, P06-1.5H-N) at 0.75 million cells per well. The following day, cells were stimulated with LPS (at a final concentration of 100 ng/ml) or H_2_O as a control.

### Library design

Perturbation libraries were constructed based on prior knowledge of inflammatory response pathways to enrich for impactful genetic perturbations. Gene targets were chosen from multiple published studies and screens, including: genome-wide CRISPR screens for TNF production in LPS-stimulated bone marrow-derived dendritic cells (423 genes at 20% FDR)^148^; Nos2 production screens in TNFα-stimulated BMDCs (454 genes at 10% FDR)^23^; phagocytosis screens in U937 cells (596 genes at 10% FDR)^24^; NFκB response screens in NALM-6 cells (263 genes at 20% FDR)^40^; LPS transcriptional regulators identified through Perturb-Seq in THP-1 cells (666 genes)^149^; E3 ligase regulators from BMDC Perturb-Seq studies (271 genes)^150^; cell morphology regulators from optical pooled screens (123 genes)^151^; and 417 manually curated genes, yielding a union of 3,090 genes. The sgRNA sequences were designed based on crisprVerse^152^. The library was optimized by iteratively replacing guide RNAs to ensure a minimum pairwise Hamming distance of 3 within the first 15 nucleotides of spacers, while guides containing excessive homopolymers (>10 cytosines) were excluded. Overall, for optical pooled screens, 13,042 sgRNAs were chosen to target 3,050 genes with up to four guide replicates, 824 sgRNAs targeting 103 olfactory receptor genes, and 40 non-targeting sgRNAs. For Perturb-Seq, 4,454 sgRNAs were selected to target 1,030 genes with up to 4 guide replicates, along with 300 sgRNAs targeting 75 olfactory receptor genes and 40 non-targeting guides as controls.

### Library cloning

The PerturbView vector was derived from CROPseq-puro-v2, with the U6 promoter replaced by a chimeric U6/T7 promoter (PerturbView-puro-v2). The oligo pool was synthesized by Twist Bioscience (San Francisco, CA), amplified by dial-out PCR^153^ using Ex Premier DNA Polymerase (Takara Bio, RR370A), cloned into the PerturbView vector with Golden Gate cloning using BsmBI (New England Biolabs, E1602S), transformed into Stellar electrocompetent cells (Takara Bio, 636765) and packaged into a lentiviral library while maintaining >1,000× coverage at every step.

### Lentiviral library preparation

HEK293T cells were seeded into a 15cm dish (Corning, 430599) at a density of 24 million cells per plate in Opti-MEM with GlutaMAX supplement (Invitrogen, 31985088) supplemented with 5% FCS, 1 mM sodium pyruvate (Fisher Scientific, 11360070) and 1x MEM nonessential amino acids (Fisher Scientific, 11140050). The following day, cells were transfected with the expression plasmid, psPAX2 and pMD2.G at a molar ratio of (1:2.8:2.4) using Lipofectamine 3000 (Thermo Fisher Scientific, L3000015). Media was refreshed with ViralBoost Reagent (ALSTEM, VB100) after 6 hours. Viral supernatant was collected 1 day after transfection and filtered through a 0.45-μm filter. The viral supernatant was further concentrated by adding a Lenti-X concentrator (Takara, 31231) and centrifuging at 4°C for 45 minutes. The concentrated supernatant in OPTI-MEM was kept at -80°C before use. Lentiviral titer was quantified before the screens using a CellTiter-Glo kit (Promega, G7570). For all experiments, an MOI of <0.2 was used.

### Cell harvesting for Perturb-Seq

After 9 h of stimulation, cells were incubated with AccuMAX at 4°C for 15 min and at room temperature for 5 min, and then further incubated with Accutase at 4°C for 15 min for dissociation. Cells were then washed with PBS and resuspended in 50μL PBS+ 2% BSA. Cells were resuspended in 100 µl of CITE staining buffer (2% BSA, 0.01% Tween-20 in PBS) and 5 µl of TruStain FcX blocking antibodies (Biolegend, 101319) were added, followed by incubation for 10 min on ice. Cells were again pelleted and resuspended in 100 µl of CITE staining buffer and 5 µl of CITE-seq antibody pool (∼1:500 final dilution for each antibody) (Biolegend, 199901) were added to each sample, followed by incubation for 30 min on ice^6^. Cells were then stained with 7 BioLegend hashing antibodies (Biolegend 155811, 155815, 155817, 155823, 155825, 155827, 155829) as previously described^25,154^. H2O-treated cells were spiked into the LPS-stimulated samples to comprise 1.67% of the total population (aiming for 1,500 out of 85,000 cells). 85,000 cells were loaded per channel on 15 10x GEM-X Single Cell 3’ channels according to the manufacturer’s instructions^155^.

### PerturbView Protocol

Cells were fixed at 9 hours post-stimulation in 4% PFA in PBS for 30 minutes, followed by three washes with PBST and permeabilization with 0.2% Triton X-100 for 10 minutes. For immunofluorescence, cells were blocked with 3% BSA in PBS for 45 minutes. Sequential immunofluorescence was performed using the IBEX method^156^ with the antibody panels listed in **Table S5**. For each staining round, cells were incubated with antibodies for 3 hours at 37°C, washed three times with PBST, post-fixed with 4% PFA for 5 minutes, and washed another three times with PBST. Cells were incubated in 200ng/mL DAPI for 10 min, then in 50ng/mL during imaging. After imaging, fluorophores were bleached by incubation with freshly prepared 1 mg/ml lithium borohydride (Strem, 50-901-13696) in water for 30 minutes, followed by three washes with PBST. *In situ* sequencing (ISS) was then performed starting from the ethanol permeabilization step^14^.

### Perturb-Seq data preprocessing

To assess gene expression levels, sequencing reads were aligned to the mm10 reference genomes using CellRanger (v8.0.1). Surface protein tag abundances, hashtag antibody barcode abundances and Celltag viral barcode library reads were estimated using the cumulus_feature_barcoding (v0.11.3)^157,158^.

### Perturb-Seq data processing

#### Initial quality filters

Cells that had less than one UMI barcode in any of the available modalities (HTO, gRNA, ADT, or gene expression) were filtered first, followed by removing cells with high (> 10%) mitochondrial UMI counts, low RNA UMI counts (<3.5k) or low protein counts (<100).

#### Perturbation assignment

To assign cells to hashtags, HTO counts were de-multiplexed with demuxEM, using a minimum UMI HTO count of 10. Cells were similarly assigned to perturbations using the guide RNA barcodes, with a minimum UMI gRNA (or gRNA + T7) count of 2.

#### Genomic features filtering

Genes with less than 500 UMIs were removed and then only the top 10,000 variable genes (identified using Scanpy ^159^) were retained. Transcripts for mitochondrial proteins and ribosomal proteins were filtered as well to reduce potential confounding effects in the downstream analysis.

#### Low-quality cluster removal

Perturbation-agnostic cell clusters were computed as follows. For this analysis only, data was normalized using scanpy’s normalize_total function, followed by logarithmic transformation. Cells were clustered using the leiden algorithm (resolution 0.5) after principal component analysis (50 components). Out of the 12 distinct clusters, one cluster of 3,494 cells (∼1% of the total number of cells in the dataset) with particularly low UMI counts was not enriched for any specific perturbations. It was therefore identified as a low-quality cell cluster and removed from subsequent analyses.

#### Guide effect analysis and filtering

First, perturbations with less than 30 cells per guide were first filtered out. Then, TotalVI was applied ^160^ to the whole dataset and the energy distance^161^ between perturbed cells and control cells, for each guide perturbation was computed. A p-value was calculated using an empirical null formed from the control cells. Specifically, the null hypothesis was calculated from 10,000 samples of 1,000 control cells each. A Benjamini-Hochberg False Discovery Rate (FDR) was calculated. Only gene perturbations with at least two guides with consistent effects (out of four) were retained, following previous work^150^. To assess consistency, guides that were not significant were first removed from the analysis. Then, an embedding of the pseudobulk profile of the cells with each guide was computed, and its similarity to the other guide’s pseudobulk profile embedding was assessed by computing the Spearman correlation coefficient of the embeddings between each pair of guides pseudobulk profiles. Gene perturbations for which at least a pair of consistent guides (Spearman ρ⩾ 0.2) could not be identified were removed from the analysis.

#### Description of MoCAVI

MoCAVI (Multi-Omic Contrastive Analysis with Variational Inference)^35^ is a contrastive latent variable model designed to disentangle perturbation-induced variation from background cellular heterogeneity in single-cell multimodal data. It takes as input single-cell measurements across one or more modalities (e.g., gene expression and surface protein abundance), along with perturbation assignments, batch covariates, and a partition of cells into a baseline (control) environment and multiple perturbation-specific environments. MoCAVI learns two complementary latent spaces: a background space that captures sources of variation present in control cells, and a salient space that captures environment-specific variation attributable to perturbations while controlling for background effects. The model outputs cell-level embeddings in both spaces, which can be used for visualization, perturbation effect quantification, and downstream statistical analyses.

#### Data preparation for MoCAVI

One important consideration before applying MoCAVI is the composition of the background dataset, and the specification of each environment. The background dataset is usually composed of the control cells. In this manuscript, MoCAVI was applied using the cells with a perturbation targeting olfactory receptor genes as background samples. Each environment from the target dataset was then specified as all the cells with a specific gene perturbation, for each perturbation. For MoCAVI analysis specifically, perturbations with non-significant effect must be filtered out in order to ensure disentanglement is successful. This analysis was therefore applied specifically on the perturbations that passed the above filtering method.

#### MoCAVI analysis of PerturbCITE-seq data

MoCAVI^35^ was applied on the gene expression and surface protein profiles. The batch variable was set to be the library identifiers. The model was trained with 15 salient variables, and 10 background variables.

#### Modality-specific MoCAVI embeddings

In MoCAVI, variational distributions from each modality and each type of latent variable (salient / background) are probabilistically combined using a product of expert formulation. We compute an RNA-only salient embedding of the cells by passing their gene expression through the expression-specific salient encoder, and keeping only the mean estimate of the modality-specific variational distribution.

#### Background Analysis

To interpret the background latent variables, Hotspot analysis^36^ was performed on the control cells, as embedded by MoCAVI’s background space, to identify clusters of genes that covary in the latent space. Using only the control cells for this task is important, as it ensures that Hotspot does not use any data related to the perturbation to construct this molecular map. After clustering of the gene-gene correlation matrix from Hotspot, gene sets were annotated using Metascape^162^. Additionally, the score of each of the Hotspot background gene programs was calculated for every cell gene expression profile (including perturbed cells) with a z-score normalization, using the control cells as reference.

#### Embedding calculation for salient space

Cell-specific embeddings were calculated using MoCAVI’s salient space and corrected by Mahalanobis whitening where the covariance matrix was computed on the control cells.

### Preprocessing for PerturbView data

Images were processed using the SCALLOPS (Scalable Library for Optical Pooled Screens) software^163^. Phenotypic images were acquired using ×20 magnification for resolution, and ISS was acquired at ×10 magnification with 2×2 binning for speed For each cycle of phenotyping and ISS images, flatfield and darkfield were estimated by median and minimum values for each pixel location, respectively. After illumination correction, for each well and cycle, images were stitched together using ASHLAR^164^ to represent a whole-well image. Images from later cycles were registered to the first cycle using DAPI channels by affine transformation followed by B-spline transformation using ITKelastix^165,166^ based on wsireg parameters^167^. Nuclei were segmented based on a DAPI channel in the first cycle of phenotyping images using Stardist^168^. A cytoplasmic foreground mask was generated from a composite (per-pixel sum) of phalloidin, Vimentin, Tomm20, and Iba1 channels to represent cytoplasm; the composite was adaptively thresholded when a pixel is 1% brighter than its counterpart in a Gaussian-blurred image (σ=30). Labeled nuclei then served as seeds for Voronoi-based propagation within the cytoplasmic foreground to obtain whole-cell boundaries^169^. To prepare segmentation masks for ISS, segmentation was transformed based on a registration from a DAPI channel of phenotyping images to ISS. Spot detection, base calling and phenotypic feature extraction were conducted as described previously^14,163^.

Cells with no assigned barcodes and cells that were assigned to two barcodes such that the intensity of the first barcode is not twice as bright as the second barcode were removed.

Median intensity was calculated for nucleus, cytoplasm, and whole cell for each assigned cell and channel. Similarly, the minimum, maximum, and 25th and 75th percentile were calculated for each channel, as was the correlation across channels for each cell, resulting in 1,960 features in total.

### PerturbView data processing

#### Feature selection

Features that could introduce outliers in the cell embeddings (and therefore constitute noise in downstream analysis) were removed if their interquartile range or standard deviation was abnormal across all features (below 0.2 after logarithmic transformation). Additionally, features were excluded if more than 0.5% of cells exhibited expression values exceeding 5 standard deviations from the mean, resulting in 1,155 features per cell.

#### Cell outlier filtering

Outlier cells were defined based on individual interpretable features and removed as follows: (**1**) Cells with the bottom 5% or top 0.5% of brightness, or bottom and top 2% length or bottom and top 2% area, (**2**) Cells with bottom 0.5% or top 0.5% of brightness correlation between DAPI and other channels (indicating potential registration issues). Then, cells scored as the top 3% outliers from each well based on IsolationForest scores were removed. During this step, we ensured that no more than 3% of the cells were removed per perturbation by capping the fraction of cells that could be removed per perturbation to exactly 3%. This avoided the error of removing entire perturbations that could appear as outliers with respect to the whole population.

#### Control guide outlier analysis

Outlier control guides were defined by generating a pseudobulk profile from the cell profiles with each control guide, computing a principal component analysis with 50 components from the pseudobulk profiles, and identifying outliers with three methods: isolation forest, elliptic envelope and local outlier factor, all from scikit-learn^170^. Control guides predicted as outliers by at least two methods were removed from the analysis.

#### Embedding calculation

Cell-specific embeddings were calculated from measurements on selected features as follows. Positive features were log-transformed, then normalized with respect to control perturbations taking into account each cell’s location on the plate, by subtracting, for each feature in each cell, the mean measurements of its 200 closest control cells in the well, and dividing by the interquantile range of that feature in the well. Principal component analysis was performed with 70 components on those normalized values. Finally, embeddings were obtained by Mahalanobis whitening where the covariance matrix was computed on the control cells, for each well separately.

#### Differential abundance analysis

The median value was computed for each marker across the cell profiles with each perturbation. A p-value was calculated using an empirical null formed from 100,000 samples of 1,000 control cells each, and a Benjamini-Hochberg FDR was calculated.

#### Guide effect analysis and filtering

Perturbations with less than 150 cells per guide were filtered out. Based on single-cell embeddings, an energy-distance between the perturbed cell profiles and control cell profiles was calculated for each guide. A p-value was calculated using an empirical null from 10,000 samples from 1,000 control cells each, and a Benjamini-Hochberg FDR was calculated. To assess consistency, guides that were not significant were removed, an embedding of the pseudo-bulk profiles of cells carrying each guide was computed, and the similarity between each pair of guides was assessed by the Spearman correlation coefficient (ρ) of their pseudobulk profile embeddings. The largest subset of guides that had ρ ≳ 0.2 was retained. Gene perturbations for which at least a pair of consistent guides could not be identified was removed from the analysis, retaining 5,638 gRNAs (1,576 genes).

### The EB-MoCAVI generative model

For perturbation *r* ∈ {*ctrl*, 1 …, *R*}, we denote by *e*_*r*_ the embedding of the phenotype derived from OPS data. In this notation, *ctrl* denotes the control condition, and *R* denotes the number of distinct significant perturbations or regulators. Let *n* ∈ {1 …, *N*} be a cell, where N is the total number of cells in the dataset.

For cell *n*, let *x*_*n*_ denote the *G*-dimensional vector of RNA counts (*G* genes) and *y*_*n*_ the *P*-dimensional vector of protein measurements (*P* proteins). Let *b*_*n*_ be a *B*-dimensional one-hot vector describing the batch index in case the data presents batch effects. Let *t*_*n*_ ∈ {*ctrl*, 1 …, *0*} be its assigned perturbation label as a categorical variable. We assume *0* ≤ *R*, which means that the set of perturbations for which embeddings are available is a superset of all the perturbations present in the Perturb-Seq data. This allows us to impute gene expression for any of the *R* perturbations for which OPS embeddings are available. By abuse of notation, we introduce the cell-level variable 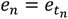 to denote the embedding of perturbation *t*_*n*_ for cell *n*. This means that cells with the same perturbation have the same value of *e*_*n*_. Finally, we use ; to refer to all the parameters of the generative model.

EB-MoCAVI estimates a generative distribution *p*_*θ*_(*x*_*n*_, *y*_*n*_ ∣ *b*_*n*_, *t*_*n*_), whose mathematical expression depends on whether cell *n* is a control cell (*t*_*n*_ = 0) or it was exposed to a genetic perturbation (*t*_*n*_ ≠ 0). We first specify the perturbed case, and discuss how our treatment of control cells differs below.

#### Empirical priors

We posit that a background cell representation 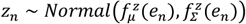 encodes the heterogeneity of phenotypes in control cells. Similarly, the salient cell representation 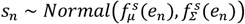 encodes the effect of genetic perturbations. In both cases, the neural networks 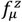,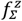,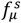 and 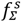 represent how each perturbation can change each of the latent distributions for every perturbation *t*_*n*_.

#### Gene expression likelihood

For every gene *g* ∈ {1, …, *G*}, the RNA count *x*_*n*g_ for gene *g* in cell *n* follows the negative binomial distribution

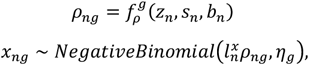

where 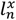 denotes the RNA size factor, and *η* denotes the gene-specific inverse dispersion parameter. *ρ*_*n*_ is the output of a neural network that takes as input *z*_*n*_, *S*_*n*_ and *b*_*n*_, and applies a softmax non-linearity to the output to ensure that the output can be interpreted as a gene expression frequency. The RNA size factor is estimated from data as 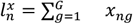, while *η* is treated as a parameter to be inferred via maximum likelihood. This follows closely the framework of scVI ^62^ and lvm-DE ^171^.

#### Protein abundance likelihood

For every protein *p* ∈ {1, …, *P*}, the protein abundance *y*_*np*_ for protein *p* in cell *n* follows a mixture of negative binomial distribution, defined as follows:

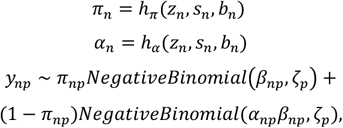

where the mixture weight *π*_*n*_ and the fold-change of the foreground protein abundance *α*_*n*_ are outputs of neural networks *h*_*π*_ and *h*_*α*_, that both take as input *z*_*n*_, *S*_*n*_ and *b*_*n*_. ζ_*p*_ is the inverse-dispersion parameter of the negative binomial distributions. The background protein abundance *β*_*n*_ is learned via (amortized) maximum likelihood. *h*_*π*_ has a sigmoid non-linearity for its output layer, *h*_*β*_ and has an exponential non-linearity for its output layer, and *h*_*α*_ has a ReLU non-linearity with an offset of 1 for its output layer. The offset ensures identifiability of the two components of this mixture (i.e., *α*_*np*_*β*_*np*_ must be the mean of the protein abundance after denoising). This setup closely follows the one from TotalVI^160^.

#### Generative distribution for control cells

In the case of a control cell (*t*_*n*_ = 0), we simply force the value of the salient variables *S*_*n*_ to be equal to zero:

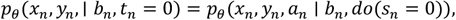

where the *do*(⋅) operator denotes a deterministic intervention, as in other contrastive analysis work^31,33,34^.

#### Inference for EB-MoCAVI

The model evidence *p*_*θ*_(*x*_*n*_, *y*_*n*_ ∣ *b*_*n*_, *t*_*n*_) cannot be computed because of the intractable integral over the latent variables. Therefore, exact Bayesian computations are impossible, and we resort to variational inference to approximate the posterior distributions, as well as to learn the model parameters including those for the empirical prior^172^.

#### Variational distribution

The variational distribution takes the following parameterization:

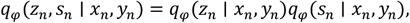

where each distribution is modeled with a product of expert distributions ^173^ to incorporate information from each modality, for example:

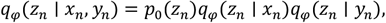

where *p*_4_ is the isotropic Gaussian distribution, and each of the variational distributions *q*_*φ*_(*z*_*n*_ ∣⋅) is a Gaussian with diagonal covariance. The product of experts of Gaussian distributions is Gaussian, with mean and covariance matrices available in closed-form. The parameters for both of those distributions are the output of neural networks (encoder networks) taking a single modality as input. We proceed similarly for *S*_*n*_:

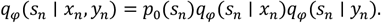

We proceed similarly for the control cells, but solely relying on the distribution *q*_*φ*_(*z*_*n*_ ∣ *x*_*n*_, *y*_*n*_), since *S*_*n*_ is deterministically set to zero.

#### Evidence lower bound

The evidence lower bound (ELBO) for a perturbed cell *n* is defined as:

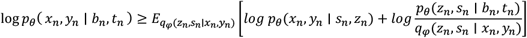

Derivations appear in **Supplementary Note 5**.

#### Iterative training procedure

Direct optimization of the ELBO with the empirical priors specified above is challenging because the neural network priors can fail to properly disentangle the background and salient representations from the observed data, an undesired behavior we have seen in practice.

We therefore developed a two-stage iterative training procedure that ensures proper disentanglement, while leveraging the empirical priors for improved generalization. We first train EB-MoCAVI using non-informative priors *z*_*n*_ ∼ *Normal*(0, *I*_*p*_), 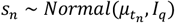 to establish proper disentanglement, where *μ* ∈ *R*^0×*q*^ represents how each perturbation shifts the salient distribution. This ensures that the variational distributions *q*_*φ*_(*z*_*n*_ ∣ *x*_*n*_, *y*_*n*_) and *q*_*φ*_(*S*_*n*_ ∣ *x*_*n*_, *y*_*n*_) learn to properly separate the background cellular heterogeneity from perturbation-specific effects.

Second, we freeze all parameters from the likelihood as well as the variational distributions and optimize the empirical prior networks 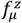,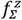,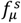,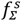 using the ELBO. This is a natural procedure, as it amounts to the maximum likelihood estimation of the empirical prior parameters using samples drawn from the learned variational distributions.

#### Implementation *details*

We optimize the ELBO relying on stochastic gradient estimates, evaluated by sampling 128 observations from each data set, as well as samples from the variational distribution. We update the model parameters with the Adam optimizer with weight decay ^174^. We use a standard multi-layer perceptron architecture with common activation functions (for example, exponential, softmax, sigmoid) to encode the variational and generative distributions.

#### Salient encoder regularization

As highlighted in previous implementations of deep generative models for contrastive analysis^33,34,175^, latent spaces *z*_*n*_ and *S*_*n*_ may contain redundant information when the number of latent variables *p* or *q* increases. To avoid this, we regularize the encoder networks parameterizing the approximate posterior over the salient space, following Multi-GroupVI^34^ and ContrastiveVI^31^. More precisely, let us denote the approximate posterior over the salient space as

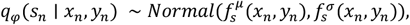

where 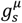 and 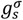 designate the mean and variance of the approximate posterior as a function of the input data. We add to the composite ELBO the following penalization:

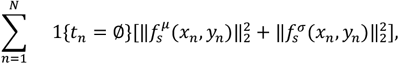

which corresponds to the squared Wasserstein-2 distance between the approximate posterior and the Dirac at zero (i.e., what the value of *S*_*n*_ should be for background cells).

#### Predictive distributions of the EB-MoCAVI model

The key component for deriving predictive distributions for EB-MoCAVI consists of methodology for sampling latent variables from a query perturbation *t*^*^. For imputation purposes, we assume that we only have the embedding 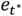, whereas for denoising purposes, we also have access to some observations 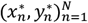 from that perturbation. We seek to approximate the posterior 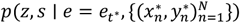 that combines information from the prior 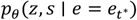 and the observations 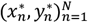, using our Bayesian framework. Using Bayes’ rule and assuming a tight variational bound, we obtain:

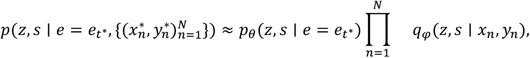

which is tractable as a product of experts of Gaussian distributions. Interestingly, we have found this principled solution to yield poor results, likely because of drastically different scale of magnitudes in the variances from the prior (pseudobulk-level) and each cell’s posterior (single-cell level).

We therefore developed an adaptive weighting scheme that interpolates between prior only (pure prediction) and single-cell posteriors (pure imputation) based on the available sample size. This approach is motivated by empirical Bayes shrinkage theory (introduced by Robbins et al.^134^. See Maritz and Lwin^176^, for a comprehensive treatment), which shows that combining estimates from multiple sources can improve overall performance through shrinkage toward a population mean. For a perturbation *t*^*^ with *N* observed cells, we use the following mixture distribution as an approximate posterior:

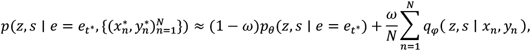

where *ω* is an adaptive weight that depends on the sample size *N* and allows the model to smoothly transition from prior-dominated inference (when few cells are available) to data-dominated inference (when many cells are observed).

The predictive distribution for the latent variables can then be used for downstream tasks, such as differential abundance analysis, using the methodology from lvm-DE^171^.

For completeness, we describe here the case of differential expression analysis, comparing two perturbations. For gene *g*, and pair of cells *α, β*, we wish to compute the probability of differential abundance (here, expression). First, we define differential expression as a probabilistic event:

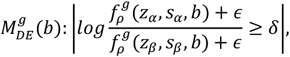

where g designates a small offset used for stability, as in, and i is the desired level of effect size. We then marginalize the latent variables, as well as categorical covariates to calculate the posterior probability of differential expression:

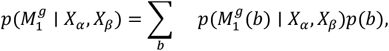

where each individual posterior probability 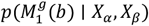 is approximated by plugging-in the approximate posterior. For example, *p*(*z*_*α*_, 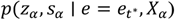, *X*_*α*_) for cell *α*.

In order to compare two cell populations *A* = (*α*_1_, …, *α*_*A*_) and *B* = (*β*_1_, …, *β*_*B*_), we average the posterior probabilities of differential abundance for random pairs of cells in each population. If one group of cells is part of the control cells (*t* = 0), then we replace *S* by 0 in all of the above calculations.

The calculation of the posterior probabilities allow for either the calculation of a Bayes factor, or binary labels that indicate the genes that are differentially abundant with a controlled posterior expected false discovery rate at a desired level^171^.

#### Differences between MoCAVI and EB-MoCAVI

Our empirical Bayesian version of MoCAVI departs from the original MoCAVI method in two ways: (1) The prior for MoCAVI’s latent variables has a neural parameterization taking as input the embeddings from OPS data, instead of being a free variable to optimize during inference. The training procedure has to be altered to accommodate for this change; and (2) New sampling subroutines must be implemented to access the posterior predictive distribution and impute both latent embeddings and differential expression of unseen perturbations, as well as denoised perturbations.

#### Filtering of highly-impacted genes, and gene program inference

Each dataset originally captured 10,000 highly-variable genes (HVGs), as identified by Scanpy during the data pre-processing. MoCAVI was run on all 10,000 HVGs, including the differential expression analysis. We used the DE analysis to keep only genes that were changed in at least ten gene perturbations (FDR < 0.20 and |LFC| > 0.1). In the primary screen, the top 4,000 most variable genes (out of the 4,612 significant genes) were kept. We referred to those as the highly-impacted genes.

Inference of the fine gene programs was performed by clustering the genes using the log-fold changes estimates from MoCAVI across all perturbations. Clustering was completed with the Leiden algorithm, and 55 clusters were identified. Hierarchical clustering was then performed to merge those clusters into 15 coarse clusters.

#### Gene program-protein correlation

To quantify the relationship between gene programs and imaging phenotypes, the regulator–burden correlation framework of Ota et al.^86^ was adapted to imaging readouts. For each gene program, significant perturbations were identified by comparing the mean log-fold change (LFC) of its constituent genes with a null distribution generated from 10,000 randomly sampled gene sets matched for program size. The correlation between perturbation effects on each gene program and its corresponding imaging feature was computed using only perturbations with a significant effect on either the gene program or the imaging feature (FDR < 0.05). Gene programs with fewer than 10 significant perturbations were excluded from the analysis.

#### Transcriptional target enrichment analysis

Gene sets for RELA transcription factor targets (ENCODE Transcription Factor Targets) were obtained from Harmonize 3.0 and converted to mouse orthologs^59,60,177^. Statistical significance was assessed using permutation tests, where null distributions were generated by randomly sampling gene sets of equivalent size from the 4,000 most variable genes (n=50,000 permutations). Empirical p-values were calculated as the proportion of permuted samples with an overlap count as or more extreme than the observed overlap (two-sided test), with a pseudocount added to avoid zero p-values. Multiple hypothesis testing was addressed using the Benjamini-Hochberg FDR procedure. FDR of <0.2 was used to call significance, and signed significance scores were calculated as the sign of the deviation multiplied by FDR.

#### Benchmarking of unseen perturbation effect imputation

The ability of several methods to predict expression responses for unseen perturbations was assessed, using methods selected to reflect different types of prior information (OPS, text embeddings from Gene Cards, gene pathways) and computational strategies (deep generative models, linear models, graph-neural networks). Each method was assessed for its accuracy in imputing log-fold-changes for perturbations held out during training.

#### Performance metrics

#### Mean squared error (MSE) on top-*k* differentially expressed genes

The MSE between predicted and observed log fold changes was computed for the top most differentially expressed genes. For each perturbation *p*, let *y*_*p*_ ∈ *R*^*G*^ and 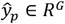 denote the ground truth and predicted log fold change vectors across *G* genes, respectively. The set 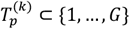 of indices corresponding to the top-*k* most differentially expressed genes was identified based on absolute log fold changes in the ground truth and the MSE Top-*k* for perturbation *p* was defined as:

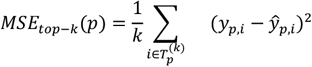

*k* = 20 was used to focus on genes with the strongest perturbation effects. The overall MSE Top-*k* metric was defined as the average across all test perturbations:

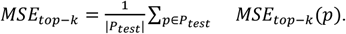

#### Area under the precision-recall curve (AUPRC)

To evaluate the ability to identify differentially expressed genes, gene selection was treated as a binary classification task. For each perturbation *p*, genes were defined as differentially expressed if their absolute log fold change exceeded a threshold *τ* = 0.5: 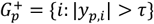. The absolute predicted log fold changes 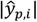 were used as confidence scores and precision and recall curves were computed across all genes and perturbations. The AUPRC was calculated by flattening predictions and ground truth labels across all test perturbations and computing:

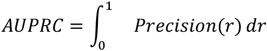

where the precision at recall level *r* is computed from the concatenated predictions and binary labels across all test perturbations.

#### k-nearest neighbors Jaccard similarity

To assess if a method correctly captures similarity between perturbations k-nearest neighbor structures were compared between predicted and observed perturbation spaces. For each test perturbation *p*, its *k* nearest neighbors in both the ground truth log fold change space and the predicted space were computed using Euclidean distance:

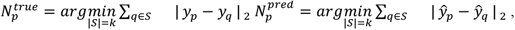

and the Jaccard similarity between the true and predicted neighbor sets was computed as:

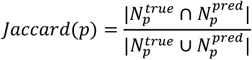

*k* = 50 was used and overall metric was defined as the average Jaccard similarity across all test perturbations.

##### Competing Methods

EB-MoCAVI was compared to the following methods for perturbation effect imputation:

### Random

A baseline method that randomly shuffles observed log fold change values to assess the performance floor.

### GEARS

A graph neural network-based method that uses gene pathways as prior knowledge to predict perturbation effects^66^. GEARS models gene-gene interactions through knowledge graphs and can impute expression profiles for unseen perturbations based on pathway relationships.

### Linear

A regularized linear regression baseline that predicts pseudobulk expression levels from perturbation embeddings, followed by log fold change computation relative to control conditions. Ridge regression with cross-validation was used to select optimal regularization parameters.

### Perturbation embeddings

The linear model is evaluated separately using each of three distinct types of perturbation embeddings as input features. (1) OPS derived embeddings derived. (2) Gene embeddings from pre-trained language models that capture semantic relationships between gene names and functional annotations (GenePT)^178^. (3) Embeddings derived from the genome-wide perturbation screen dataset from K562 cells^64^.

#### Data splitting strategy

Two data splitting strategies were employed to evaluate generalization performance. Random splitting consisted of a random assignment of 20% of gene-level perturbations to the test set, ensuring independent and identically distributed train-test splits. As a more challenging evaluation, pathway splitting consisted of *k*-means clustering the log fold change perturbation profiles into *k* = 20 clusters, and then holding out entire clusters for testing to reach approximately 20% test size. Pathway splitting tests generalization to perturbations with novel mechanisms of action by ensuring that test perturbations come from different functional clusters than training perturbations.

#### Benchmarking of perturbation effect denoising

To evaluate EB-MoCAVI’s ability to improve perturbation effect estimates when limited single-cell data is available, cells from test perturbations were systematically subsampled, and the ability to recover accurate perturbation effects by incorporating empirical priors was assessed by comparing the log-fold change estimates from the method (that has access to only the subsampled cell profiles) to the observed LFCs on the original dataset.

#### Experimental Design

A random 80/20 train-test split of perturbations was employed, and the fraction of cells retained for test perturbations was varied, (1%, 5%, 10% and 20%) while keeping all cells for the training perturbations and controls.

#### Baseline comparisons

As no established approaches exist for this task (to the best of our knowledge), three plausible approaches were devised within the EB-MoCAVI framework: (**1**) pure prediction, in which the few cells available are discarded, and only OPS data are used to predict the effect of the (now unseen) perturbation; (**2**) pure imputation, in which the cells from sub-sampled perturbations are included in the MoCAVI analysis, and differential expression analysis is performed using the standard methodology (ignoring the prior); and (**3**) adaptive shrinkage, in which the cells from sub-sampled perturbations are included in the EB-MoCAVI analysis, and the shrinkage estimator is used to combine the prior and the sparsely available data into an estimate of cell states.

#### Evaluation of denoising performance using protein-protein interaction (PPI) data

Ground-truth biological neighborhoods were defined using the PPI databases STRING^38^ and CORUM^37^, filtered to include only genes whose perturbations are present in our dataset and requiring minimum connectivity (≥3 neighbors). For each candidate perturbation and embedding method, the *k*-nearest neighbors (*k* systematically varied from 5 to 50) were identified using cosine similarity and coverage scores were calculated as the fraction of PPI interactors from the PPI data that are recovered among the top-*k* most similar perturbations.

### Human genetics data processing

#### Quantitative trait analysis

For quantitative traits including monocyte, neutrophil, lymphocyte, and eosinophil counts, denoised rare-variant burden statistics from Beckman et al.^89^ were used, processed through GeneBayes^90^ a flexible platform that can improve the estimation of many gene-level properties, such as rare-variant burden scores, by incorporating a population genetics model with machine learning on gene features to reduce noise in burden effect estimates.

#### Disease trait analysis

For binary disease traits including Type 2 diabetes (Study Accession: GCST90085502, GCST90085503, GCST90085512, GCST90083728), inflammatory bowel diseases (GCST90081491), ulcerative colitis (GCST90081490, GCST90084220), asthma (GCST90085447), COPD (GCST90085455), and rheumatoid arthritis (GCST90084378, GCST90085490), rare-variant burden statistics were obtained from the exome sequencing analysis of 454,787 UK Biobank participants in Beckman et al.^89^. Loss-of-function burden tests were retained using variants with minor allele frequency < 0.1%. Effect sizes were calculated as log_10_(odds ratio).

#### Gene identifier mapping

Human gene identifiers were mapped to mouse orthologs using the Mouse Genome Informatics homology database^177^. When multiple human genes mapped to the same mouse gene, the variant with the strongest burden effect was retained to avoid redundancy.

#### PROGRAM burden effect calculation

For each gene program *π*, the program burden effect *γ*_*π*_ was calculated as the mean effect size *γ*_g_ of all genes *g* within the program on the trait of interest. Statistical significance was assessed using permutation tests, where null distributions were generated by randomly sampling 1,000 gene sets of equivalent size from the pool of all program genes with mapped mouse orthologs. P-values were calculated as the proportion of permuted samples with effects equal to or more extreme than the observed program effect, with a pseudocount added to avoid zero p-values. A Benjamini-Hochberg false discovery rate (FDR < 0.1 for quantitative traits, FDR < 0.25 for binary traits) was calculated across programs within each trait.

For binary traits, prevalence-adaptive minor allele count (MAC) filters were applied (MAC ≥ 25, 50, or 150 for traits with estimated prevalence > 5%, 1.5–5%, or < 1.5%, respectively) to ensure sufficient expected case carriers per gene. Trait prevalence was estimated from allele counts and burden test standard errors. Gene programs with fewer than 20 genes after MAC filtering or fewer than two significant disease associations were excluded from analysis.

#### Regulator-Burden Correlation Analysis

The regulator-burden correlation *ρ*_*π*_ statistics was introduced by Ota et al.^86^ and measures the correlation between genetic burden effects and perturbation effects within each program, controlling for selective inference bias. For each gene program *π*, our analysis was focused on a set of regulators *r* that significantly affect the trait (*γ*_*r*_ significant for p < 0.05) or the program (*β*_*r*→*π*_ significant for FDR < 0.05). To assess the significance of *β*_*r*→*π*_, the mean LFC of the program’s genes was compared to a null distribution derived from 1,000 randomly sampled gene sets of matching size, with the Benjamini-Hochberg FDR calculated for the empirical p-values (FDR < 0.05). *ρ*_*π*_ was then computed as the partial correlation between burden effect sizes *γ*_*r*_ and program-level effect *β*_*r*→*π*_, controlling for *S*_*het*_ values (which represent the selective constraint on heterozygous loss-of-function variants as estimated by GeneBayes) using ordinary least squares regression. This metric provides improved constraint estimates especially for shorter genes. FDR was then calculated across programs within each trait.

#### Selection of targets and guides for secondary (validation) Perturb-Seq

260 genetic perturbations were selected for the secondary screen (**Table S7**) as follows. 193 perturbations were selected with an algorithmic procedure from 1,082 imputed profiles, by imputing the salient MoCAVI embedding for each perturbation, then applying Leiden clustering at resolution 2.0 (47 clusters), removing any cluster with no imputed genes or in the lowest 25^th^ percentile of energy distance (estimated from salient embedding), and, for each remaining cluster, randomly selecting up to 4 genes already profiled in the main screen (total 103 perturbations) and 4 unseen genes (total 90 perturbations). Another 67 perturbations were selected as follows: 24 perturbations with low cell count in perturb-seq but detected as hits in OPS (only) to study denoising (out of those 24, only 11 had enough cells for downstream analysis), and 20 of the nearest neighbors of the Srp proteins. For guide selection, the single sgRNA with the highest OPS energy distance for each target was included, along with 28 guides with the lowest energy distances for the Olfr controls. Non-targeting controls were not included in the library.

---

**Figure S1.**
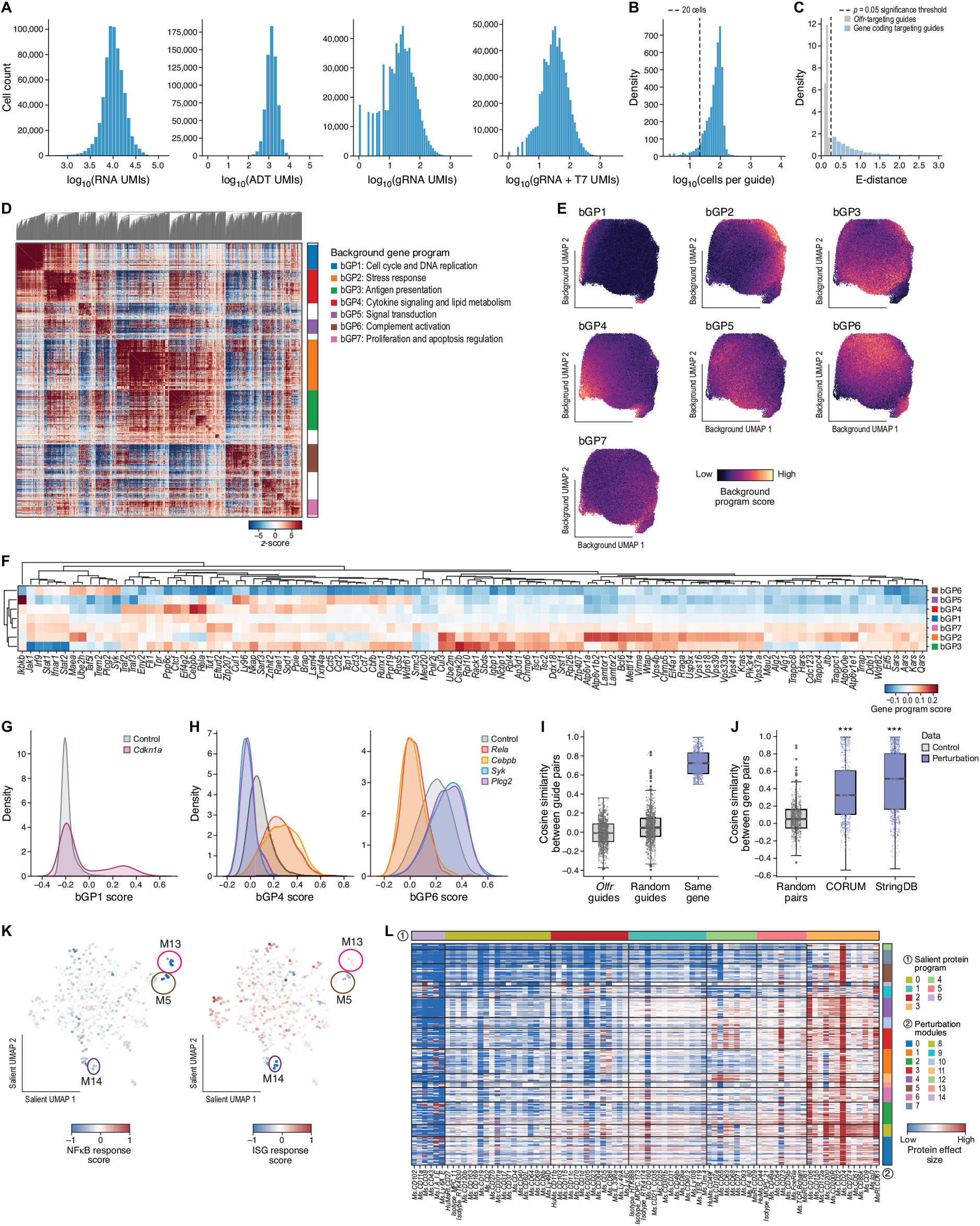
related to Figure 2. Perturb-Seq analysis and the MoCAVI background space. (A-B) Quality control metrics. Distributions of number of gene expression UMIs per cell (A, left, x axis), number of ADT UMIs per cell (A, second from left, x axis), number of gRNA UMIs per cell (A, second from right, x axis), number of gRNA + T7 UMIs per cell (A, right, x axis), and number of cells per guide after filtering (B) in the Perturb-Seq experiment. (C) Impactful perturbations. Distribution of energy distances (x axis) of individual guide perturbations for targeting guides (blue) and guides targeting olfactory receptor genes (gray). Dashed line: significance threshold (P<0.05, empirical p-value). (D-F) D, Background gene programs (bGPs). D, Auto-correlation statistics (by Hotspot ^36^, z-score, color bar) between genes (rows, columns) according to their expression weighted by distances in MoCAVI’s background space and ordered by hierarchical clustering (top) and labeled by background gene program membership (color legend, right). E, UMAP of MoCAVI’s background representation of the Perturb-Seq profiles (dots) colored by score (color bar) of each background gene program. F, Scores (color bar) for each bGP (rows) in the pseudobulk profiles (columns) for the perturbations with the strongest impact in the background space. (G-H) G, Select regulatory relations with background programs. G, Distributions of bGP1 scores (x axis) in profiles for cells carrying a guide targeting Cdkn1a (red) or control guides (blue). H, Distributions of bGP4 (left) and bGP6 (right) scores (x axis) in profiles for cells carrying a guide targeting Plcg2 (purple), Rela (red), Cebpb (orange), Syk (blue), or control guides (gray). (I-J) Consistency of the salient space. Distribution of cosine similarity scores (y axis) between pairs of pseudobulk profiles of guides targeting the same gene, two controls (Olfr) or random pairs of genes (I, x axis) or between pairs of pseudobulk profiles of perturbations of genes encoding proteins in the same pathway (by CORUM or STRING) or random pairs of genes (x axis, J). (K) MoCAVI representation of the salient latent space. UMAP cell profiles (dots) in MoCAVI’s salient space, colored by gene signature scores of NF-kB response (left) or ISG response (right) gene program. (L) Effect on CITE-Seq protein markers. Log-fold changes (color bar) of expression of select CITE-seq protein markers (columns, ordered by protein program (bar 1)) across genetic perturbations (rows, ordered by module (bar 2)).

**Figure S2.**
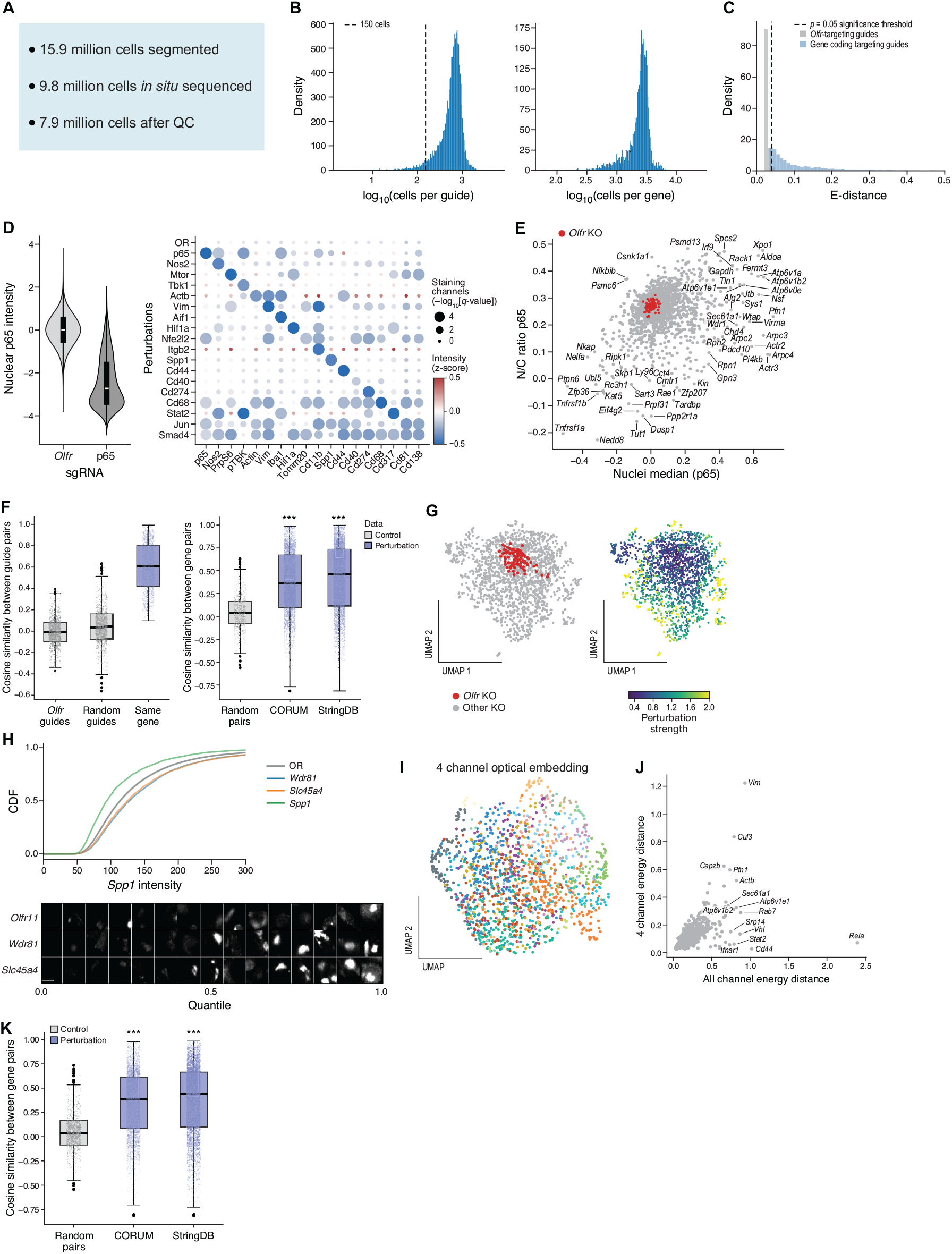
related to Figure 2. OPS analysis. **(A) Overview of PerturbView filtering**. (B) Quality controls. Distributions of number of cells with each guide (left) and with guides targeting each gene (right). (C) Impactful perturbations. Distribution of energy distances (x axis) of individual guide perturbations for targeting guides (blue) and guides targeting olfactory receptor genes (gray). Dashed line: significance threshold (P<0.05, empirical p-value) (D) Left: Distribution of nuclear p65 protein staining intensity (y axis) in cells with guides targeting Rela (the gene encoding p65) or controls (x axis). Right: Relative intensity vs. cells with control guides (dot color, z-score) and FDR (dot size, -log_10_(q-value); one-sided adjusted empirical p-values) of expression of proteins (columns) that are either encoded by genes targeted by perturbations or are the established indirect protein targets of perturbed genes (rows). (E) Regulators impacting p65. Median intensity in the nucleus (x axis) and nucleus to cytoplasm intensity ratio (y axis) of p65 in cells with guides targeting each gene (gray dots) or Olfr controls (red dots). (F) Distribution of cosine similarity scores between pseudobulk profiles of pairs of guides targeting the same gene, controls (Olfr), or targeting random gene pairs (left, x axis), or pairs of pseudobulk profiles of perturbations of genes encoding proteins that are members of the same pathway (by CORUM or STRING) or of random gene pairs (x axis, right). (G) OPS perturbation profile embedding. UMAP embedding of the pseudo-bulk profiles of each gene perturbation (dot), colored by control vs. target set (left) or by energy distance (right). (H-I) Biological impacts highlighted by OPS. H, Top, cumulative distribution functions (CDF) of Spp1 intensity (x axis) for cells with guides targeting Spp1 (green), Wdr81 (blue), Slc45a4 (orange), or olfactory receptors (OR, controls; gray line; shading standard deviation across olfactory receptor genes). Bottom, representative Spp1 images under each perturbation (rows) for cells sampled from each of the 15 intensity quantiles (according to the mean cellular Spp1 intensity for each perturbation) (columns, left to right) . (I-K) Four morphological channels are highly informative. I. UMAP embedding of pseudo-bulk 4-channel optical feature profiles of each perturbation (gene level, dots), colored by OPS perturbation modules from Figure 2H. J, Energy distance (x and y axes) between perturbed and control cells for different genes (dots) from all imaging channels (x axis) or just four morphological (Cell Painting) channels (y axis). K, Distribution of cosine similarity scores between pairs of pseudobulk profiles (four morphological channels only) of cells perturbed for genes encoding proteins that are members of the same pathway (by CORUM or STRING) or between random gene pairs (x axis, right).

**Figure S3.**
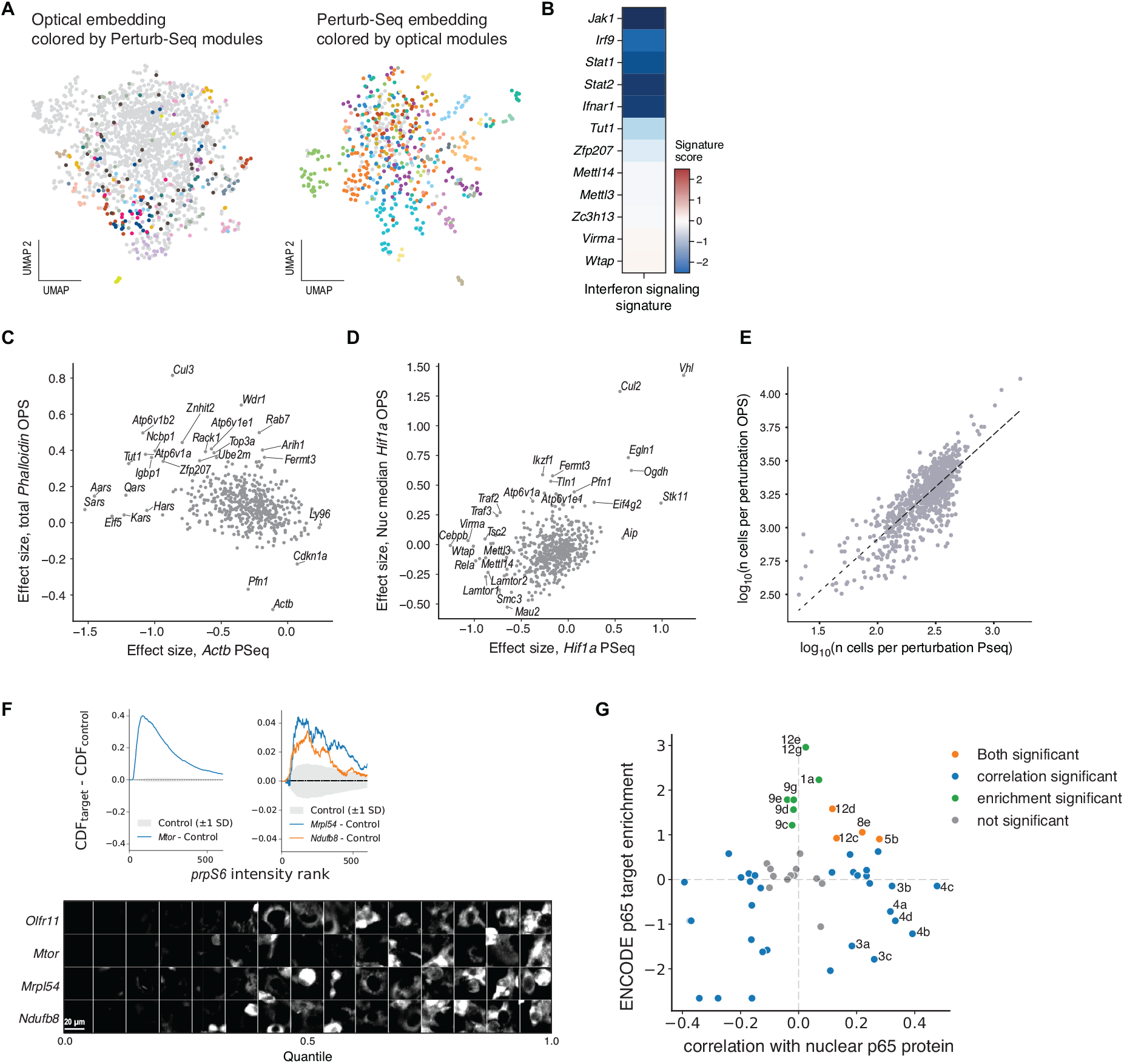
related to Figure 3. Global concordance and finer distinctions between OPS and Perturb-Seq. (A) UMAP embedding colored by perturbation modules of different modalities. Left, optical UMAP embedding colored by transcriptional perturbation modules. Right, transcriptional salient UMAP embedding colored by OPS modules. Perturbation modules are described according to colors in Figure 2. (B) m6A component perturbations do not affect interferon gene programs. Signature score (color bar) of an interferon signal program in pseudobulk Perturb-Seq profiles of different gene perturbations (rows). (C-D) Differential expression effect size (x axis, from Perturb-Seq) and protein staining (y axis, from OPS) for Actb (C) and Hif1a (D) for different gene perturbations (dots). (E) Number of cells profiled per gene perturbation (dot) for all jointly profiled perturbations across each assay (OPS, y axis and Perturb-Seq, x axis). (F) Top: Delta cumulative distribution functions (CDF) of prpS6 intensity for cells targeting Mtor (left) and Mrpl54, Ndufb8 (right) relative to the mean of olfactory receptor controls (Control; gray; shading standard deviation across control genes). Bottom: representative prpS6 images under each perturbation (rows) of cells sampled from each of 15 intensity quantiles (according to the mean cellular prpS6 intensity) (columns, left to right). (G) Pearson’s r between nuclear median p65 intensity and gene program score (x axis) and enrichment or depletion of p65 transcriptional targets (y axis, signed log_10_(FDR) based on empirical p-values) of each gene program, colored by significance on Pearson’s r (blue), p65 target enrichment/depletion (green) or both (orange).

**Figure S4.**
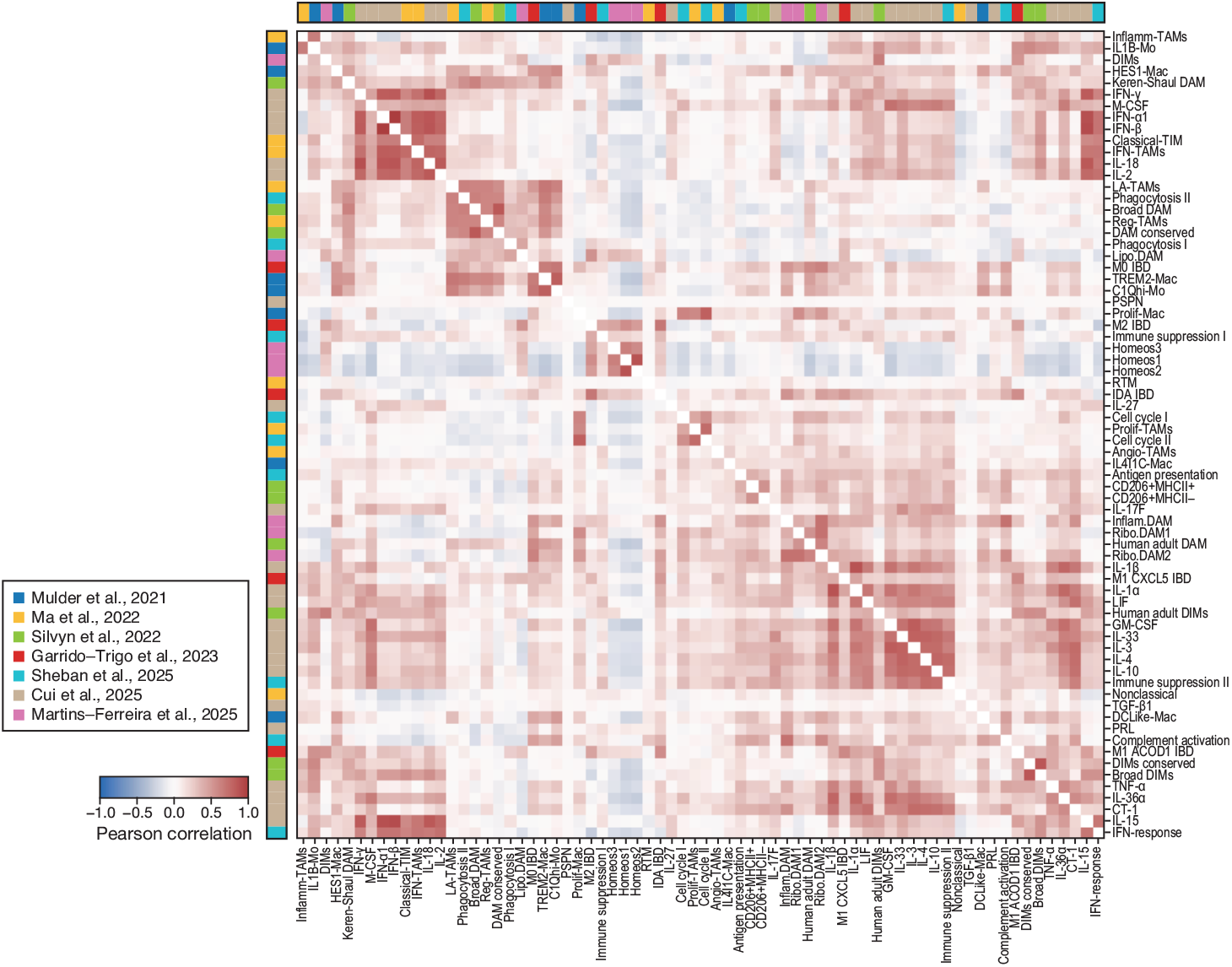
related to Figure 5. Unperturbed (control) cells are less informative for macrophage gene programs. Pairwise Pearson’s correlation coefficient (color bar) between curated gene signatures (rows, columns; ordered as in Figure 5G) based on their scores across measured and imputed perturbation profiles. Bars on left and top: source studies.

**Figure S5.**
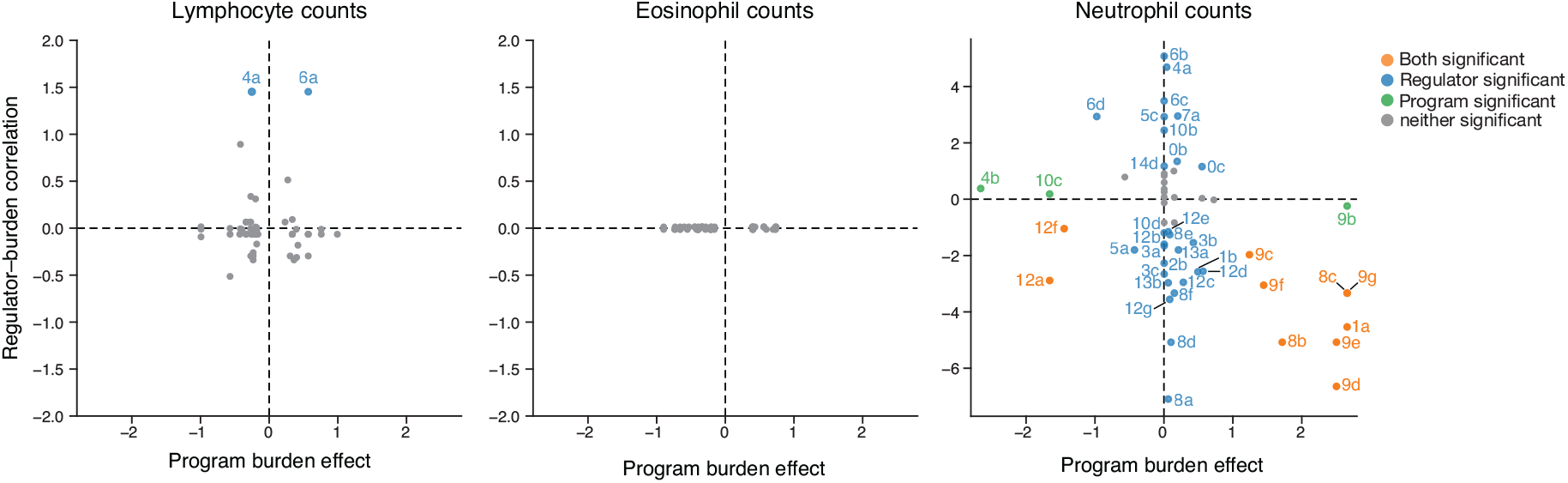
related to Figure 6. Regulators and programs associated with blood cell counts. Program burden effects (signed log_10_(FDR), empirical p-value, x-axis) and regulator-burden correlation (signed log_10_(FDR), partial Pearson correlation, y-axis) of 55 sub-programs (dots) in lymphocyte (left), eosinophil (middle) and neutrophil (right) cell counts traits from the UK Biobank, colored by significant (FDR<0.1) associations with program burden effects (orange), regulator burden correlation (blue) or both (green).

**Figure S6.**
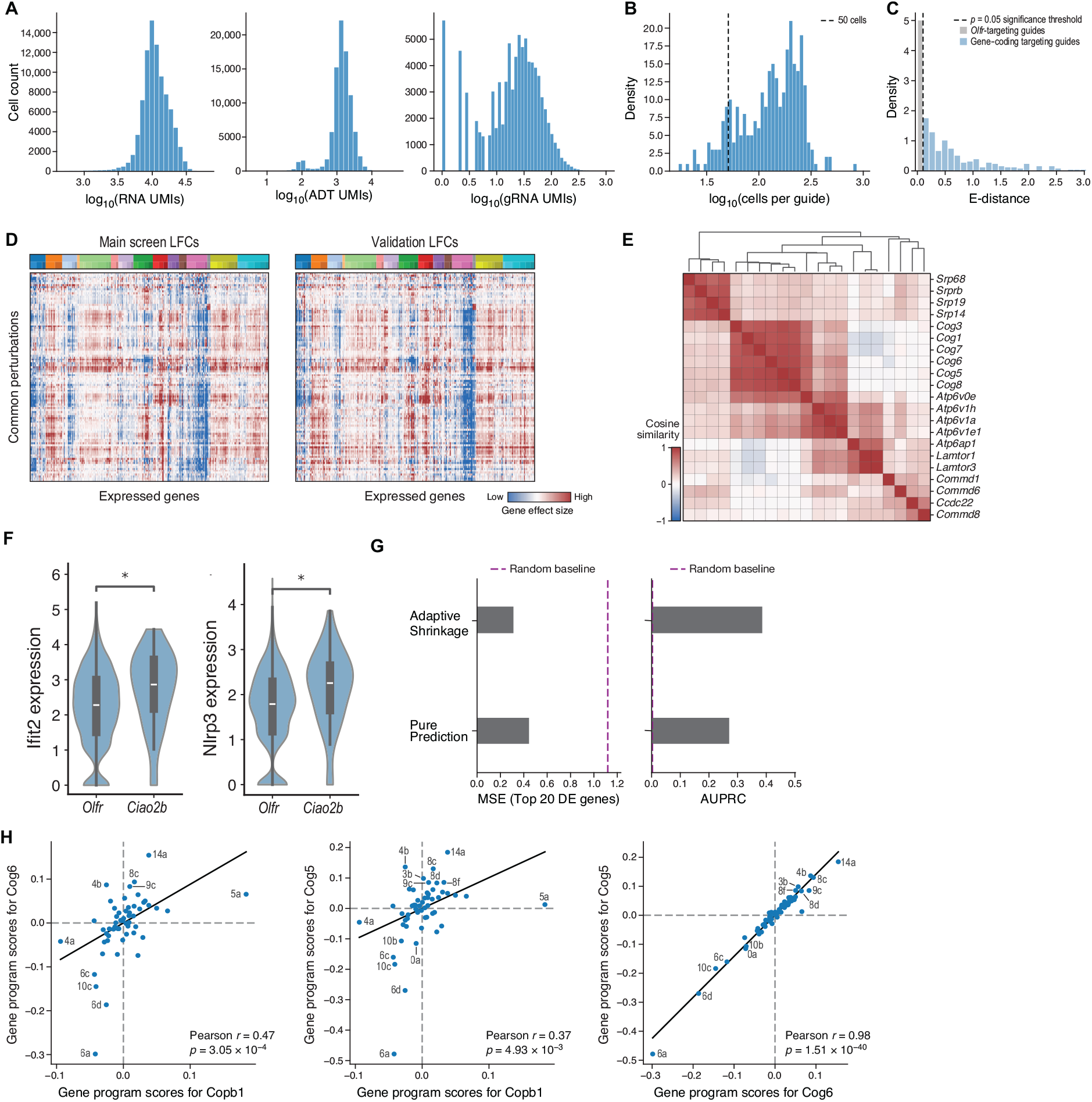
related to Figure 7. Validation of EB-MoCAVI’s imputation and denoising by a secondary Perturb-Seq screen. (A-B) Quality control metrics. Distributions of the number of UMIs per cell (A, left, x axis), number of ADT UMIs per cell (A, middle, x axis), number of gRNA UMIs per cell (A, right, x axis), and number of cells per guide after filtering (B) in the Perturb-Seq experiment. (C) Impactful perturbations. Distribution of energy distances (x axis) of individual guide perturbations for targeting guides (blue) and guides targeting olfactory receptor genes (gray). Dashed line: significance threshold (P<0.05, empirical p-value). (D) Qualitative evaluation of reproducibility. Log-fold changes of genes (columns) from salient gene programs introduced in Figure 2 (color labels on top) in the primary (main) screen (left) and the secondary (validation) screen (right) in each of 143 pseudobulk perturbation profiles measured in both screens (rows). (E) Cosine similarity in PCA embedding (color bar) of pairs of pseudobulk profiles of cells perturbed for SRP components and their neighbors (rows, columns). (F) Distribution of normalized expression levels (y axis) of Ifit2 (left) and Nlrp3 (right) in cells with guides targeting Ciao2b or Olfr controls (x axis). (G) Quantitative evaluation of the denoising. MSE (top 20 genes, x axis, left) and AUPRC (x axis, right) of expression profiles denoised by adaptive shrinkage or using the pure prediction approach across 11 perturbations with sparse data in the primary screen to their measurements in the secondary screen. Results are reported as the average across five random seeds. (H) Validation of imputation-based predictions. Log-fold score changes (x and y axes) of salient gene programs (dots) in pseudobulk profiles of cells perturbed for Copb1, Cog6 or Cog5 in the validation screen.

