## Supplementary Information for "Scalable multimodal mapping of macrophage regulatory architecture by integrating optical and transcriptomic pooled screens"

**Supplementary Note 1. Perturb-Seq Preliminary Data Analysis**

For Perturb-Seq, we loaded 85,000 cells per lane onto 15 10x GEM-X lanes, using cell hashing for overloading and protein quantification, and performed scRNA-seq. As a non-stimulated control, we included a pool of cells (~2% of the population) treated with $H_2O$. We obtained profiles from 654,184 non-empty droplets containing 510,024 single cells and 144,160 multiplets, followed by dedicated PCR to detect the guide RNA in each cell. We also utilized the PerturbView promoter to perform linear amplification of barcodes and obtain additional counts for the guide RNA readout (**Methods**). Using the combined counts, we observed a marginal improvement of sgRNA capture of 7,345 cells compared to the dedicated PCR solely, resulting in a final count of 334,633 single cells assigned to a unique perturbation (**Methods**). This resulted in a high-quality dataset with a median number of 10,230 UMIs from gene expression and 1,371 UMIs from the protein abundance (**Figure S1A**), with a median number of 74 cells per guide (311 cells per gene perturbed, **Figure S1B**).

We then proceeded to characterize the effect of genetic perturbations. As a quality control step, we performed outlier analysis to filter out control perturbations that had an unintended effect on the cells (**Methods**). We then filtered guides that either had no effect, or had potential off-target effects. After this filtering step, we retained 2,191 guide perturbations targeting 625 unique genes. In order to account for potential off-target effects, we then retained only those perturbations that had at least two significant guides with agreeing effects (assessed using the Spearman correlation coefficient on the pseudobulk profiles for each guide, as previously described [1], retaining 2,119 guides targeting 547 genes for downstream analyses.

**Supplementary Note 2. Biological interpretation of Perturb-Seq background gene programs**

To better delineate the heterogeneity in macrophage activation, we applied Hotspot [2] to identify gene programs that define distinct cellular states within the background space. This analysis revealed seven background gene programs (bGPs) representing key cellular processes involved in macrophage responses to LPS stimulation (**Figure 2B; Figure S1D,E**). bGP1 was enriched for genes involved in cell cycle progression and proliferation (including Cdk1, Ccnb1, Ccnb2, Plk1, Mcm2-10, Cenpa, Cenpe, Cenph, Cenpf, and DNA repair components Rad51, Rad51b, Brca1, Brca2), suggesting enhanced proliferative activity in a subset of macrophages. bGP2 captured early inflammatory responses with genes associated with cytokine production (Il12b, Cxcl1, Cxcl2, Cxcl3), cellular stress (Atf3, Gadd45b, Ddit3), and redox balance (Prdx1, Txnrd1, Srxn1, Nqo1). bGP3 was characterized by interferon-stimulated genes (Ifit2, Ifit3, Rsad2), antiviral response components (Gbp4, Gbp5, Nos2), and immune activation markers (Cd69, Itgax). bGP4 encompassed alternative activation markers including lipid metabolism genes (Pparg, Lpl, Cav1), extracellular matrix components (Col4a1, Col4a2, Col4a5), and anti-inflammatory signaling. bGP5 included genes from signal transduction and immune receptor pathways (complement receptors C3ar1, C5ar1, Tlr8, transcriptional regulators Irf8, Maf). bGP6 was defined by complement system activation (C1qa, C1qb, C1qc, Cfb), inflammatory cytokines (Il1b, Il1a), and classical activation markers (Nlrp3, Lcn2). Finally, bGP7 captured stress response and cell survival pathways, including cell cycle arrest genes (Cdkn1a) and apoptosis regulators (Prkn, Pdcd4).

Scored genetically perturbed cells according to their expression of these background programs showed that genetic knockouts could produce distinct phenotypic effects that aligned with their known molecular functions (**Figure S1F**). For instance, cells with Cdkn1a knockout displayed the strongest enhancement of bGP1 (**Figure S1G**), indicating increased proliferative activity consistent with p21's role as a critical cell cycle checkpoint protein. In addition to such expected cell cycle effects [3], we observed a striking opposing contradiction between the regulation of bGP4 (alternative activation/metabolic) and bGP6 (complement/classical inflammation), revealing fundamental trade-offs in macrophage activation strategies. Core inflammatory transcription factors showed inverse effects on these programs: Rela and Cebpb knockout enhanced bGP4 while suppressing bGP6 (**Figure S1H**). Conversely, knockout of immunoreceptor signaling components Syk and Plcg2 reduced bGP4 while enhancing bGP6. This aligns with their established roles: Syk mediates immunoreceptor tyrosine-based activation motif (ITAM) signaling that can promote anti-inflammatory responses [4,5], while PLCγ2 participates in lipid signaling cascades essential for metabolic reprogramming in alternatively activated macrophages [6–8]. Overall, this coordinated opposition demonstrates that macrophages face resource allocation constraints, choosing between classical complement-mediated inflammatory responses versus metabolically demanding alternative activation programs characterized by enhanced lipid metabolism and tissue remodeling functions.

**Supplementary Note 3. Biological analysis of Perturb-Seq salient gene programs and modules**

We investigated the perturbation effects based on MoCAVI's salient space. Leiden clustering of the single cell data in the salient space revealed 15 co-functional modules of perturbation (**Figure 2B/C, bottom**). These modules covered many aspects of the regulatory networks governing macrophage function and inflammatory responses, including core inflammatory signaling pathways (NF-κB, MAPK, mTOR signaling; module 0), transcriptional machinery and chromatin remodeling (RNA polymerase II complex, histone modifications; module 1, 6, 11), RNA processing and splicing regulation (spliceosome components; module 3, 10), protein synthesis and quality control (ribosomal proteins, ER stress response, aminoacyl-tRNA synthetases; clusters 4, 5), vesicular trafficking and lysosomal function (V-ATPase complex, TRAPP complex; clusters 7, 8), LPS sensing and TLR4 signaling (TLR4, MyD88, IRAK4; module 13), type I interferon responses (JAK-STAT pathway components; module 14), molecular chaperones (CCT complex; module 12), and epigenetic regulation including m6A RNA methylation and cohesin complex components (module 9). This comprehensive coverage demonstrates how genetic perturbations can systematically be dissected into the modular regulatory architecture underlying macrophage activation and inflammatory responses.

As an initial characterization of these modules, we scored the pseudobulk profile associated with each perturbation with a data-driven, 300-gene signature of LPS response, defined by applying differential expression analysis to non-stimulated cells versus unperturbed cells exposed to LPS (**Table S8**). Three of the modules impaired the LPS response most severely: module 13 (TLR4/MyD88 signaling), module 14 (JAK-STAT pathway), and module 5 (N-glycosylation machinery). We then scored each of the three modules for their effects on well-established gene signatures of NF-κB targets (*Tnf*, *Il1b*, *Il6*, *Nfkbia;* the immediate inflammatory response triggered by TLR4 recognition of LPS) and interferon-stimulated genes (ISGs) (*Ifit1*, *Mx1*, *Gbp* family, capturing the secondary amplification phase driven by autocrine and paracrine cytokine signaling). These signatures allowed us to determine whether perturbations disrupt specific regulatory pathways or broadly impair macrophage activation (**Figure S1K)**.

Module 13 perturbations (including *Tlr4*, *Myd88*, *Irak4*) showed the most severe reduction in NF-κB targets, consistent with their role in signal initiation - the primary pathway converting LPS recognition into transcriptional activation. Module 14 perturbations (*Jak1*, *Jak2*, *Stat1*, *Stat3*) specifically abolished ISG expression, reflecting their function in signal amplification through cytokine signaling cascades that sustain and broaden the immune response. Module 5 perturbations, encompassing N-glycosylation machinery (*Alg1*, *Alg2*, *Ddost*, *Rpn1*, *Rpn2*) and protein quality control systems, showed a distinctive pattern: significant reduction in NF-κB responses with minimal effects on ISG expression. This pattern reflects the critical role of protein glycosylation in receptor function - N-glycosylation of Tlr4 and Md-2 is essential for proper LPS recognition and receptor assembly, making glycosylation perturbations phenocopy direct disruption of the MyD88 pathway while preserving alternative interferon production routes. These results reveal that robust macrophage activation requires coordinated regulation across multiple layers: transcriptional initiation (MyD88), cytokine-mediated amplification (JAK-STAT), and receptor assembly and quality control (N-glycosylation machinery). Each layer contributes unique regulatory mechanisms, and disruption at any level impairs the overall immune response through distinct molecular signatures

To comprehensively characterize how the 15 perturbation modules regulate macrophage functional states, we systematically analyzed their effects across each of 15 salient gene programs (defined via clustering of the perturbation effect sizes estimated by MoCAVI; **Methods**), revealing distinct regulatory logic for each module (**Figure 2D**). The 15 gene programs captured different aspects of macrophage biology: inflammatory signaling (GP0: including Il6, Il10, Nfkbia, Socs3, and Ptgs2), oxidative stress (GP1: including Bach1, Txnip, Tnfaip3, Sesn3, and Ephx1), interferon / antiviral (GP2: including Ifit1, Ddx60, Oas1b and Mx1), membrane trafficking (GP3: including Rabac1, Snx7, Stx3, and Optn), redox / autophagy (GP4: including Cat, Prdx1, Txnrd1, Slc7a11 and Gabarapl1), immune signaling (GP5: including Tnf, Il12b, Tlr7, Nfkb2, and Pik3cg), phagocytic activation (GP6: including Itgam, Lyz2, Fcgr1, and Trem1), glycolysis/hypoxia (GP7: including Hk2, Pfkp, Vegfa, and Egln1), metabolic / trafficking (GP8: including Gclc, Pik3ca, Lamp2, and Arfgef1), chromatin regulation (GP9: including Kdm6a, Setd1b, Atrx, Eed, and Sin3a), cytoskeleton / mitochondria (GP10: including Actn1, Tubb6, Idh2 and Park7), antigen presentation (GP11: including H2-Aa, H2-Eb1, H2-DMb1 and Cd74), ribosome biogenesis (GP12: including Nop56, Nop58, Nop10, Fbl, and Ncl), immune regulation / cytoskeleton (GP13: including Ptpn22, Il21r Inf2 and Septin8), and metabolic / UPR (GP14: including Xbp1, Hspa5, Herpud1, and Trib3).

We performed a parallel analysis of the surface protein features profiled in our Perturb-CITE-Seq experiment. We clustered perturbation effect sizes for these protein features in the same manner as for gene expression, identifying six salient protein programs that summarize how perturbations reshape macrophage surface state. These protein programs captured complementary axes of macrophage biology, including an adhesion/tetraspanin and activation module (PP0; including CD9, CD44, ICAM1/CD54, and CD73), an activated myeloid/APC program with inhibitory checkpointing (PP1; including CD64, CD40, CD11c, and PD-L1/CD274, alongside CD200–CD200R), an antigen-uptake and myeloid lineage program (PP2; including CSF1R/CD115, DEC205/CD205, CD1d, and CD301a), a checkpoint/immune-interaction axis (PP3; including VISTA/VSIR, PD-1/CD279, and PVR/CD155), a phagolysosomal/vesicle–lysosome program (PP4; including CD68, CD63, and CD107a/LAMP1), and a professional APC maturation program (PP5; including MHC-II I-A/I-E, CD86, and PIR-A/B). Together, these protein programs provide a complementary surface-level readout to the 15 gene salient programs, enabling us to connect perturbation-driven transcriptional rewiring to concrete shifts in macrophage surface identity and immune function.

These systematic regulatory relationships demonstrate that macrophage responses involve coordinated regulation across multiple functional axes, with specific modules controlling distinct aspects of inflammatory and metabolic programs through predictable biological mechanisms.

**Supplementary Note 4. Regulatory impacts on univariate markers in OPS**

In addition to the regulation of p65 and global analysis across features, this note highlights key regulators governing the individual markers Nos2, Iba1, Hif1a, phospho-Tbk1, and Spp1.

For Nos2 (**Figure 2G**), 749 perturbations had significant effects (FDR < 5%) with perturbation to the Nos2 gene being the second strongest negative effect (i.e., positive regulator). The positive regulators were enriched with Jak/Stat pathway genes, while Traf3 was the strongest negative regulator, indicating Nos2 is primarily regulated by a Trif-dependent pathway rather than Tlr4 signaling. Out of 31 gene perturbations that were previously identified as Nos2 regulators in IFNg-treated BMDM screen [9] and were captured in our experiments, 26 perturbations were recovered as significant by OPS.

Among positive regulators of Iba1 (**Figure 2G**), Iba1 (Aif1) itself, Ptpn6 (SHP-1), Bcl6, Cop1, Ctdspl2, and Vhl were notable; for example, Ptpn6 loss is known to amplify inflammatory and phagocytic programs [10], supporting elevated Iba1, while VHL's role in HIF-mediated macrophage activation aligns with increased Iba1 in activated cells [11]. Strong negative regulators included Ptpn11 (SHP-2), Syk, Shoc2, Fermt3, Actr3, Arpc4, Pfn1, and Dcaf7—many of which converge on cytoskeletal remodeling and ITAM/Syk–ERK signaling. Many of the hits have been associated with microglial or neuroinflammatory response, including Trem2 (one of the strong negative regulators) as well as Tyrobp, Plcg2, and Syk, where mutations were implicated in neuroinflammation/neurodegenerative diseases [12,13].

Many regulators of Hif1a intensity (**Figure 1G**) were consistent with the multi-layered post-translational control of HIF1α stability. The strongest negative regulators included Vhl, Cul2, and Egln1, which form the canonical E3 ubiquitin ligase complex for oxygen-dependent HIF1α degradation [14,15]. Other top negative regulators included the metabolic enzymes Ogdh and Dlst; these contribute to HIF1α hydroxylation and succinylation of prolyl hydroxylase (Egln1), thereby promoting degradation [16]. Positive regulators comprised Hif1a itself, Mau2, Ddb1, Gpn2, and Ccnc, implicating roles for cohesin loading, DNA damage response, nuclear transport, and cell cycle control in sustaining HIF1α levels under inflammatory conditions.

Phospho-Tbk1 represents Tbk1 signaling activity. Active Tbk1 forms oligomers and appears punctate in images (**Figure 2F**), and thus we used the maximum intensity to rank perturbations that affected pTBK. As expected, positive regulators were enriched with Jak/Stat pathway genes (**Figure 1G**). Strong negative regulators were enriched with autophagy pathway genes, such as Rb1cc1, Atg9a, and Atg101, consistent with autophagy loss boosting STING–TBK1 via two routes—mtDNA buildup that drives cGAS–STING and failed p62/SQSTM1-dependent clearance of STING that prolongs ER–Golgi signaling [17,18]. The immunoglobulin-binding scaffolding protein Igbp1 was a positive regulator and has also been characterized to be involved in the autophagy pathway [19].

Spp1 (Osteopontin) is a secreted matricellular protein with functions in immune modulation [20]. Consistent with Spp1 being a secreted protein, genes involved in endolysosomal acidification complex (v-ATPase subunits, Atp6v0e, Atp6v1a, Atp6v1e1, Atp6v1b2) and vesicular trafficking (Sec61a1, Surf4, Arfrp1, Sys1, Rab7, Nsf) were strong negative regulators of Spp1 (**Figure 1G**). Surf4 is a cargo receptor that regulates the ER export of proteins containing an amino-terminal tripeptide motif (hydrophobic–proline–hydrophobic), a motif present in Spp1 [21]. The recent proteomic analysis in HuH7 cells identified Spp1 as a potential client [22].

Finally, we characterized regulators that impact the subcellular localization of p65, quantified as the pixel-wise correlation between DAPI and p65 intensity under cytoplasmic segmentation as a readout of nuclear translocation (N/C ratio; **Figure S2E**). This approach allowed us to distinguish regulators of translocation from those that primarily affect expression levels. As expected, perturbation of the nuclear export gene Xpo1 markedly increased p65 nuclear translocation and accumulation, consistent with impaired export. Arp2/3-complex knockouts (Arpc3/2/4, Actr2/3) primarily increased total p65 with modest translocation effects, indicating the expression-driven nuclear signal. In contrast, Tardbp (TDP-43) knockout reduced the translocation score yet slightly increased nuclear p65 abundance. We interpret this as two opposing mechanisms: loss of TDP-43 reduces p65 nuclear import/activation (consistent with prior reports that TDP-43 physically interacts and modulates p65 nuclear translocation [23]), while TDP-43 deficiency simultaneously activates innate-immune pathways (dsRNA and/or cGAS–STING) that elevate basal p65 levels. Tardbp being the significant hit for phospho-Tbk1 (**Figure 1G**) further supports the activation of the innate-immune pathways.

### Supplementary Note 5. Bayesian derivations

#### Evidence Lower Bounds

We follow derivations from the literature [24–26] and include them for completeness.

#### Perturbed cell

The evidence lower bound (ELBO) for perturbed cell $n$ is derived from Jensen's inequality as follows. We start with the evidence (i.e., log marginal likelihood):

$$log\, p_\theta(x_n, y_n | b_n, t_n) = log \int p_\theta(x_n, y_n, z_n, s_n | b_n, t_n)\, dz_n\, ds_n$$

$$= log \int p_\theta(x_n, y_n | z_n, s_n) p_\theta(z_n, s_n | b_n, t_n)\, dz_n\, ds_n.$$

We then introduce the variational distribution $q_\varphi(z_n, s_n | x_n, y_n)$ as follows:

$$log\, p_\theta(x_n, y_n | b_n, t_n) = log \int \frac{p_\theta(x_n, y_n | z_n, s_n) p_\theta(z_n, s_n | b_n, t_n)}{q_\varphi(z_n, s_n | x_n, y_n)} q_\varphi(z_n, s_n | x_n, y_n)\, dz_n\, ds_n$$

$$= log\, E_{q_\varphi(z_n, s_n | x_n, y_n)}\left[\frac{p_\theta(x_n, y_n | z_n, s_n) p_\theta(z_n, s_n | b_n, t_n)}{q_\varphi(z_n, s_n | x_n, y_n)}\right].$$

Since the $log$ function is concave, we may apply Jensen's inequality:

$$log\, p_\theta(x_n, y_n | b_n, t_n) \geq E_{q_\varphi(z_n, s_n | x_n, y_n)}\left[log \frac{p_\theta(x_n, y_n | z_n, s_n) p_\theta(z_n, s_n | b_n, t_n)}{q_\varphi(z_n, s_n | x_n, y_n)}\right]$$

Expanding the logarithm of the product and reassembling,

$$log\, p_\theta(x_n, y_n | b_n, t_n) \geq E_{q_\varphi(z_n, s_n | x_n, y_n)}\left[log\, p_\theta(x_n, y_n | z_n, s_n) + log \frac{p_\theta(z_n, s_n | b_n, t_n)}{q_\varphi(z_n, s_n | x_n, y_n)}\right]$$

**Control cells**

Following the same derivations, we get the evidence lower bound for control cells:

$$log\, p_\theta(x_n, y_n | b_n, t_n = \emptyset) \geq E_{q_\varphi(z_n | x_n, y_n)}\left[log\, p_\theta(x_n, y_n | z_n, s_n = 0) + log \frac{p_\theta(z_n | b_n, t_n = \emptyset)}{q_\varphi(z_n | x_n, y_n)}\right].$$

**Equivalence between prior fitting and maximum likelihood**

We claim that fitting the ELBO after fixing the likelihood model of the generative distribution and the variational posterior is equivalent to maximum likelihood estimation of the parameters of the neural network used in the empirical prior fitted on data drawn from the variational distribution.

To see this, we first remind the reader of the expression of the evidence lower bound for cell $n$:

$$log\, p_\theta(x_n, y_n | b_n, t_n) \geq E_{q_\varphi(z_n, s_n | x_n, y_n)}\left[log\, p_{\theta_l}(x_n, y_n | z_n, s_n) + log \frac{p_{\theta_p}(z_n, s_n | b_n, t_n)}{q_\varphi(z_n, s_n | x_n, y_n)}\right].$$

where we decompose $\theta = (\theta_p, \theta_l)$. Here, $\theta_p$ and $\theta_l$ represent the parameters of the empirical prior (resp. likelihood model). Then, because we treat the $\theta_l$ and $\varphi$ as fixed, we have that:

$$arg \max_{\theta_p} log\, p_\theta(x_n, y_n | b_n, t_n) = arg \max_{\theta_p} E_{q_\varphi(z_n, s_n | x_n, y_n)} log\, p_{\theta_p}(z_n, s_n | b_n, t_n),$$

which is inherently a maximum likelihood estimation problem, in which we simulate data from the variational distribution, and learn parameters from the empirical prior that fits these data the best.
